# Mutation of the nuclear pore complex component, *aladin1*, disrupts asymmetric cell division in *Zea mays* (maize)

**DOI:** 10.1101/2020.12.16.423105

**Authors:** Norman B. Best, Charles Addo-Quaye, Bong-Suk Kim, Clifford F. Weil, Burkhard Schulz, Guri Johal, Brian P. Dilkes

**Affiliations:** USDA, Agriculture Research Service, Plant Genetics Research Unit; Columbia, MO, 65211; Department of Horticulture & Landscape Architecture; Purdue University; West Lafayette, IN, 47907; Department of Biochemistry, Purdue University; West Lafayette, IN, 47907; Natural Sciences and Mathematics, Lewis-Clark State College; Lewiston, ID, 83501; Department of Botany and Plant Pathology, Purdue University; West Lafayette, IN, 47907; Department of Agronomy, Purdue University; West Lafayette, IN, 47907; Center for Plant Biology, Purdue University; West Lafayette, IN, 47907

**Keywords:** Nuclear Pore Complex, Asymmetric cell division, Stomata, Subsidiary cells, Maize

## Abstract

The nuclear pore complex (NPC) regulates the movement of macromolecules between the nucleus and cytoplasm. Dysfunction of many components of the NPC results in human genetic diseases, including triple A syndrome (AAAS) as a result of mutations in ALADIN. Here we report a nonsense mutation in the maize ortholog, *aladin1* (*ali1-1*), at the orthologous amino acid residue of an AAAS allele from humans, alters plant stature, tassel architecture, and asymmetric divisions of subsidiary mother cells (SMCs). Crosses with the stronger nonsense allele *ali1-2* identified complex allele interactions for plant height and aberrant SMC division. RNA-seq analysis of the *ali1-1* mutant identified compensatory transcript accumulation for other NPC components as well as gene expression consequences consistent with conservation of ALADIN1 functions between humans and maize. These findings demonstrate that ALADIN1 is necessary for normal plant development, shoot architecture, and asymmetric cell division in maize.

## Introduction

The nuclear pore complex (NPC) is a multi-protein complex that is involved in regulating the movement of macromolecules into and out of the nucleus. The NPC also plays important roles in nuclear assembly during cell division (Rasala, Orjalo, Shen, Briggs, & Forbes, 2006). NPC proteins assemble in groups of eight, as spokes, around the opening of the pore to control transport between the nucleus and cytoplasm (Lin et al., 2016; Reichelt et al., 1990). Components of the NPC are conserved across many eukaryotic species, including mammals (J. M. Cronshaw, Krutchinsky, Zhang, Chait, & Matunis, 2002), plants (Tamura, Fukao, Iwamoto, Haraguchi, & Hara-Nishimura, 2010; Tamura & Hara-Nishimura, 2013), fungi (Liu, De Souza, Osmani, & Osmani, 2009; Stuwe et al., 2015), and yeast (Rout et al., 2000). Dysfunction and/or mutation of the genes encoding NPC components has drastic effects on growth, development, and survival of eukaryotic species. In humans, genetic mutations of NPC members have been linked to a number of diseases and illnesses, including different types of cancer, neurological diseases, and autoimmune diseases (Nofrini, Di Giacomo, & Mecucci, 2016; Roth & Wiermer, 2012; Sakuma & D’Angelo, 2017).

The structure and conserved components of the NPC of plants have been identified (Fiserova, Kiseleva, & Goldberg, 2009; Neumann, Jeffares, & Poole, 2006; Tamura et al., 2010; Tamura & Hara-Nishimura, 2013). Genetic studies have functionally characterized a few NPC members in plant species. The Arabidopsis NPC member, *At*MOS7/Nup88, was shown to regulate the nuclear concentrations of a subset of defense proteins, but other proteins tested were not affected in *Atmos7* mutants (Cheng et al., 2009). Additional forward and reverse genetic approaches in plant species; primarily Arabidopsis, tobacco, and *Lotus japonica*, have further characterized the function and role of NPC members in plant defense, plant symbioses, and responses to abiotic stress (Binder & Parniske, 2018; Cheng et al., 2009; Dong et al., 2006; Roth & Wiermer, 2012; Wiermer et al., 2012; Zhang & Li, 2005). To date, no member of the NPC has been functionally characterized in *Zea mays* (maize). It is unclear to what extent the specific function of NPC members can be inferred from phylogeny. In comparisons between mice and human, mutation of WD-40 repeat containing NPC member ALADIN did not produce the same phenotypic outcomes (Huebner et al., 2006). It is unclear whether such species-specific phenotypic impacts will characterize NPC functions in plants.

Here we describe the first characterization of an NPC mutant in maize and the first description of an ALADIN mutant in plants. The *aladin1-1* (*ali1-1*) mutant was identified in an ethyl methanesulfonate (EMS) screen as a short plant with unusual tassel architecture and defective subsidiary cell asymmetric division in developing stomata. The *ali1-1* mutant results in a weak allele and crosses with a null allele identified allelic dosage effects of the *ali1* gene, demonstrating that *ali1-1* was haploinsufficient for maintaining normal development especially after phase change. RNA-sequencing identified a compensatory increase in transcripts encoding other NPC members in *ali1-1*, even during juvenile growth before the defects in SMC asymmetric division were visible. These results implicate this gene in cell division control and NPC homeostasis that may also be true for humans and contribute to AAAS disease.

## Materials and Methods

### Plant material and growth conditions

In the summer of 2010, the *ali1-1* mutant was identified in a B73 background ethyl methanesulfonate (EMS) population as M3 generation plants segregating in the line 04IAB73PS007D8 (Weil, 2009). The *ali1-2* allele was created by an EMS-targeted mutagenesis approach. B73 pollen was incubated with 6.48 mM EMS (Sigma-Aldrich, St. Louis, MO) in paraffin oil (Sigma-Aldrich, St. Louis, MO) for 35 mins then squirted onto ∼205 *ali1-1*/*ali1-1* ears using a 40 mL squeezable bottle (Hobby Lobby, Oklahoma City, OK). ∼10,000 M1 seeds were screened in the field in the summer of 2015 in West Lafayette, IN for phenotypes similar to *ali1-1*. A total of 13 plants resembling *ali1-1* were identified. One of the twelve was heterozygous for the causative single nucleotide polymorphism (SNP) in *ali1-1*. This plant was backcrossed to *ali1-1*/*ali1-1* and B73 to propagate the *ali1-2* allele.

To map the *ali1* gene, an F2 population was created by crossing Mo17 with *ali1-1*/*ali1-1* pollen and self-pollinated the F1 generation. The F2 progeny was planted in the field to collect tissue from selected wildtype and *ali1-1* phenotypical plants. Wild-type and *ali1-1*/*ali1-1* shoot apical meristem (SAM) and surrounding tissue of 15 days after planting (DAP) plants were grown in the greenhouse in trays with a 2:1 mixture of peat germinating mix (Conrad Fafard Inc., Agawam, MA) to Turface MVP (Profile Products LLC, Buffalo Grove, IL). Total plant height, stomatal index, stomatal density, and percent aberrant stomata measurements of the 8^th^ leaf were collected from plants grown in the field in the summer of 2016 in West Lafayette, IN. Plant height, internode length, tassel length, and rachis length of B73 and *ali1-1* were measured on field grown plants in the summer of 2013. The +/+, +/*ali1-1*, +/*ali1-2, ali1-1*/*ali1-1, ali1-1*/*ali1-2*, and *ali1-2*/*ali1-2* plants used for total plant photographs and epidermal peel photographs were grown in the Purdue Horticulture Plant Growth Facility in the spring of 2016 and spring of 2017, respectively. The nine individual BC1F2 plants used for DNA sequencing were grown in the greenhouse in the spring of 2014. The temperature settings were 27°C (day) and 21°C (night) with a 16 h day length provided by supplemental lighting. Plants were grown in two-gallon pots in soilless media as previously described.

### Phylogenetic analysis

Phylogenetic analysis was performed by obtaining protein homolog coding sequences of GRMZM2G180205 through Phytozome version 10.2.2 (Goodstein et al., 2012) using the Inparanoid method (O’Brien, Remm, & Sonnhammer, 2005) with a Dual Affine Smith-Waterman score greater than 1,250 and sequence similarity of greater than 50%, resulting in 37 sequences from 32 taxa. Sequences were downloaded and a phylogenetic consensus tree was obtained as previously described in (Best et al., 2016). The N-terminus sequence was trimmed from species *Manihot esculenta* (cassava4_1.026236m), *Malus domestica* (MDP0000250771), and *Solanum tuberosum* (PGSC0003DMG400017881) due to incorrect start sites based upon alignment similarities.

### Sequencing and linkage analysis of ali1 locus

To determine the DNA sequence of wild-type, *ali1-1*/*ali1-1*, and *ali1-1*/*ali1-2*, and *ali1-2*/*ali1-2*, total DNA was extracted and amplified using the primers described in Table S1. PCR products were sequenced (Psomagen Inc., Maryland) using the Sanger method (Sanger, Nicklen, & Coulson, 1977). Restriction fragment length polymorphism genotyping of *ali1-1* and *ali1-2* alleles was performed using the derived Cleaved Amplified Polymorphic Sequences (dCAPS) method (Neff, Neff, Chory, & Pepper, 1998) and separated on a 3% agarose gel in Tris-Acetate-EDTA to observe size differences. For the *ali1-1* dCAPS genotyping, polymerase chain reaction (PCR) was carried out using primers DCali-1_FOR and ZmALI_REV+4119 (Table S1). The amplified DNA was incubated with the restriction endonuclease TaqI (New England Biolabs; Ipswich, MA) resulting in an uncut wild-type band of approximately 140 bp and the *ali1-1* mutant amplicon was cleaved to 115 bp (Fig. S1). For the *ali1-2* genotyping, amplification was carried out using primers DCali-2_FOR and ZmALI_REV+3243 (Table S1) and the uncut wild-type band was about 340bp and the *ali1-2* mutant band was 320 bp following cleavage by the restriction endonuclease HindIII (New England Biolabs; Ipswich, MA). Digestion of the *ali1-2* fragments was incomplete and as a result, heterozygous and homozygous *ali1-2* plants could not be distinguished using this molecular marker (data not shown). M37W was crossed as ear parent with *ali1-1*/*ali1-1* pollen. F1 plants were selfed and F2 generation *ali1-1*/*ali1-1* plants were phenotyped and tissue was collected for genotyping by the dCAPS method in the summer of 2015. Linkage of *ali1-1* to the *rp1* locus (AC152495.1_FG002) on maize chromosome 10 was determined by crossing *ali1-1*/*ali1-1* ears with *Rp1-D21*/+ pollen (backcrossed 8 generations to B73) and then backcrossing the F1 *Rp1-D21* phenotype plants with *ali1-1*/*ali1-1* as ear parents (Pryor, 1993). The BC1 F1 generation was phenotyped for both the *Rp1-D21* and *ali1-1* phenotypes in the field during the summer of 2016.

### Bulked segregant analysis and RNA sequencing

For bulked segregant analysis (Michelmore, Paran, & Kesseli, 1991) of SNP co-segregation with *ali1-1* and mRNA accumulation differences, total RNA was extracted (Eggermont, Goderis, & Broekaert, 1996) from 2 replicates of 60 wild-type and 60 *ali1-1*/*ali1-1* tassel stem punches at the location of the lowest primary tassel branch before tassel emergence. Samples were taken from F2 individuals derived by selfing the F1 progeny from a cross of Mo17 ears pollinated with *ali1-1*/*ali1-1*. Non-stranded cDNA libraries were prepared with a TruSeq RNA sample preparation kit v2 and sequenced on a HiSeq 2000 (Illumina, San Diego, CA) using SBS v3-HS reagents for paired-end sequencing (2 x 100bp). Raw reads were quality filtered to remove reads with less than 20 quality score and adapters were clipped using Trimmomatic (version 0.22) (Bolger, Lohse, & Usadel, 2014) and fastx_clipper as a part of the FASTX-Toolkit (version 0.0.13) (http://hannonlab.cshl.edu/fastx_toolkit/). Filtered reads were aligned to the B73 reference genome (version 3.30) using Bowtie2 (version 2.2.8) (Langmead & Salzberg, 2012). Total reads and alignment rate are shown in Table S2. Polymorphisms between Mo17 and B73 were used to map the causative mutation in *ali1-1* (Data table S1). Single-nucleotide polymorphism positions were identified by aligning Mo17 Illumina sequence to the B73 reference genome and calling SNPs by SAMtools (version 1.3.1) (Li et al., 2009) “mpileup” command. Only SNPs that were homozygous non-B73 reference, had reads on forward and reverse strands, and had less than 12 reads aligning to respective position were retained and used as SNPs for distinguishing sequencing reads from B73 and Mo17 chromosomes.

To map the causative mutation in *ali1-1*, B73 and Mo17 allele frequencies were determined from the *ali1-1*/*ali1-1* and wild-type samples and plotted in 100 SNP bins for each maize chromosome (Fig. S2-S6). To identify potential causative mutations for the *ali1* phenotype, SNPs that differed between the *ali1-1* pools and B73 reference that were in coding sequences were identified using SAMtools mpileup command and filtered to be homozygous non-B73 reference, G to A or C to T transitions, and have reads on both forward and reverse strands at the position. Identified SNP positions were then annotated for effect on coding sequence and protein function using SnpEff (Cingolani et al., 2012). An Arabidopsis protein BLAST database was created using BLAST (version 2.6.0+) and used to align all maize protein sequences. Best hits were used to provide additional annotations to the maize genes using the TAIR10 annotations (Lamesch et al., 2012). The E-value score cutoff was set at 10^−5^. Therefore, if the best BLASTP result had an E-value greater than 10^−5^, the result was not included for annotation and these columns are denoted with an asterisk..

To identify differential mRNA accumulation between mutant and wild-type samples (Data table S2 and S3), reads were aligned to B73 reference genome (version 3.30) using Bowtie2 (version 2.2.8) (Langmead & Salzberg, 2012). Counts tables were created in Htseq (version 0.6.1) (Anders, Pyl, & Huber, 2015) and passed to DESeq2 (R-bioconductor; version 3.3.0) (Love, Huber, & Anders, 2014). Dfferential expression between wild-type and *ali1-1* samples were assessed using the “nbinomLRT” setting with “parametric” dispersion. Additional annotation of maize differentially expressed genes was conducted by comparing Arabidopsis (version TAIR10) and maize (version 3.30) proteomes using BLASTP (Tatusova & Madden, 1999). Genes were annotated as previously described.

### *DNA sequencing of* ali1 *mutants*

DNA was extracted from nine individual *ali1-1*/*ali1-1* plants, sonicated, and were libraries constructed using the Illumina TruSeq DNA PCR-free LT Library Preparation protocol. Paired-end 100 bp reads were generated on an Illumina HiSeq 2500 using SBS v3-HS reagents. Sequencing statistics for the nine individual *ali1-1*/*ali1-1*;B73 BC1 plants are described in Table S12. The nine fastq files were concatenated together for downstream analysis. Reads were mapped to the B73 reference genome (version 3.30) using BWA (version 0.7.12) (Li & Durbin, 2009). SNPs in coding sequence between aligned reads and the reference genome were done as previously described. Data table S6 shows high quality SNPs as called by SAMtools and annotated for coding sequence effects by SnpEff.

### Quantification of stomatal phenotypes

Epidermal imprints and peels were collected from field-grown B73 (+/+), +/*ali1-1, ali1-1*/*ali1-1, ali1-1*/*ali1-2*, and *ali1-2*/*ali1-2* plants. Epidermal cell layer imprints were produced from the widest portion of the eighth leaf when the plants were at V9 stage using ethyl-2 cyanoacrylate (Scotch SuperGlue; 3M, Maplewood, MN) applied to the leaves and then pressed onto and recovered on glass microscope slides (Thermo Fisher Scientific, Waltham, MA). Imprints of the abaxial leaf surfaces were observed by light microscopy using a UNICO H606T microscope (United Products & Instruments Inc., Dayton, NJ). The percent aberrant stomata and subsidiary cells were calculated by non-repeatedly randomly selecting viewable areas at 40X magnification until total stomata observed per sample was greater than 500 (*n*=6 per genotype). Stomatal density and indices were calculated by counting all the cells in 5 randomly selected viewable areas between vasculature tissues for each imprint at 20X magnification. An average was calculated from 5 viewable areas from each of 6 biological replicates per genotype. An additional set of wild-type B73 (+/+), +/*ali1-1, ali1-1*/*ali1-1, ali1-1*/*ali1-2*, and *ali1-2*/*ali1-2* plants were grown in the greenhouse and epidermal cell layer imprints were used for quantification of aberrant stomata from the fourth leaf at V5 stage and tenth leaf at V11 stage were conducted as previously stated.

Leaf epidermal peels were performed by cutting 1-cm squares from greenhouse-grown plants and fixing the tissue in 4% formaldehyde, 50 mM KPO_4_, 5 mM EDTA, and 0.2% saponin at pH 7.0 for at least 2 h at room temperature. Leaf tissue pieces were rinsed 3 times with ddH_2_O and incubated in 100 mM Na-acetate, 1% cellulase, and 0.5% pectinase at pH 5.0 for at least 2 h at room temperature. The tissue was then rinsed with ddH2O and the abaxial leaf epidermis was peeled from the rest of the leaf and incubated in a 1:10 dilution of 0.05% TBO and 10 mM Na-acetate until evenly stained. Epidermal peels were viewed and photographed using an Olympus BX43 light microscope with Olympus DP80 camera (Olympus Corporation, Center Valley, PA) at 40X magnification.

## Results

### Characterization and identification of the ali1 mutant

Mutagenized progeny of the maize inbred B73 (Weil, 2009) were screened in the M3 generation for reduced stature phenotypes in the field. The *aladin1-1* (*ali1-1*) mutant was identified as semi-dwarf maize plants with reduced tassel length in a single M3 family. Shown in Fig. 1 are the *ali1* mutant gross morphological phenotypes. The *ali1-1* mutation was inherited as a monogenic recessive trait after backcrossing to the wild-type B73 (Fig. 1 A,B). Field grown *ali1-1* mutants were significantly shorter than wild-type plants and exhibited a 29% or 27% reduction in plant height measured to the tip of the tassel or flag leaf collar, respectively (Fig. 1 F,G). The reduction in plant height observed in *ali1-1* mutants was due to compression of the first two and last five internodes compared to wild-type siblings (Fig. 1 A, B, and H). The largest suppression of internode length was observed in the internodes above the ears, which elongate primarily after phase change has occurred (Fournier & Andrieu, 2000). Total tassel length of *ali1-1* mutants was 40% reduced compared to wild type (Fig. 1I). Tassel and stem elongation were variably affected, and some tassels did not completely emerge from the leaf whorl. The rachis length, as measured from the lowest tassel branch to the tip of the rachis, was 41% less in *ali1-1* mutants as compared to wild-type siblings and tassel internodes were shortened along the entire distal axis of the tassel in the *ali1-1* mutants (Fig. 1J). Tassel branching was also reduced in *ali1-1* mutants (Table S3). Visible phenotypes in *ali1-1* mutants primarily affected adult tissues that developed after phase change, suggesting a progressive effect on plant development resulting from the *ali1-1* mutation. Similar to field-grown plants, the *ali1-1* mutant was significantly shorter than wild type when grown in the greenhouse in spring months of 2016 (Table S3 and S4). By contrast, when grown in the greenhouse during the shorter days of winter 2015, *ali1-1* mutants were considerably taller, possibly due to suppression of the mutant phenotype by short days and/or low light quality and *ali1-1* plants were visually indistinguishable from wild-type siblings.

**Fig. 1.**
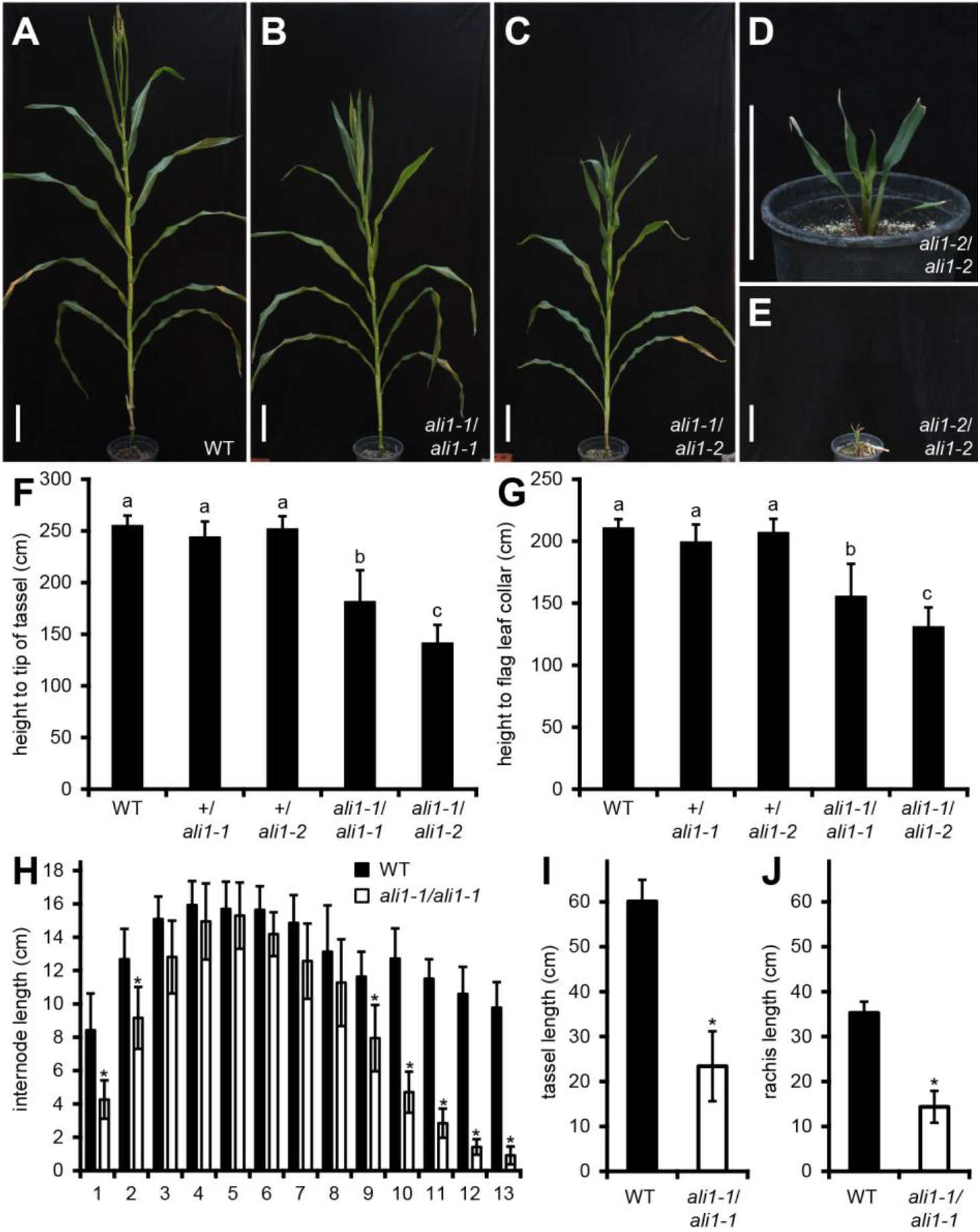
Morphological features of *ali1* mutants. (**A**) Mature wild-type (+/*ali1-1*), (**B**) *ali1-1*/ *ali1-1*, (**C**) *ali1-1*/ *ali1-2*, (**D** and **E**), and *ali1-2*/*ali1-2* plants grown in the greenhouse. (**F**) Total plant height measured to tip of tassel and (**G**) plant height measured to flag leaf collar at maturity of wild-type (*n*=17), +/*ali1-1* (*n*=16), +/*ali1-2* (*n*=20), *ali1-1*/*ali1-1* (*n*=15), and *ali1-1*/*ali1-2* (*n*=14) grown in the field. (**H**) Internode lengths of field-grown wild-type (*n*=25) and *ali1-1*/*ali1-1* (*n*=17) plants. Internode one indicates the internode closest to the soil. (I) Tassel length, including the peduncle and rachis, of wild-type (*n*=25) and *ali1-1*/*ali1-1* (*n*=17) plants. (**J**) Rachis length of wild-type (*n*=25) and *ali1-1*/*ali1-1* (*n*=17) plants measured from the lowest tassel branch to the tip of the rachis. (**A-E**) Scale bars represents 20cm. (**F-G**) Connecting letter report determined by Student’s T-test with Bonferroni corrected *p*-value (*p*<0.005) for multiple testing. (**H-J**) asterisks indicate Student’s T-test (*p*<0.01) for comparisons between wild-type and *ali1-1/ali1-1* at given internode. (**F-J**) Error bars are standard deviations.

The genetic map position for *ali1-1* was determined by bulked-segregant analysis of RNA-Seq data from F2 mutant and wild-type pools of *ali1-1* x Mo17. Reads were mapped to the B73 reference sequence and the allele frequencies at SNP positions between Mo17 and B73 (Data table S1) were scored in 100-SNP bins and graphed (Fig. S2-S5). A region near the telomere of chromosome 10S approached homozygosity for B73 in the mutant pools (Fig. S6; Data table S4-S7). To obtain additional linkage information, *ali1-1* was crossed to *Rp1-D21*/*+*, a hyper active allele of a NUCLEOTIDE-BINDING LEUCINE-RICH REPEAT (NLR) gene for resistance to *Puccina sorghi* on 10S (Pryor, 1993), and F1 progeny were backcrossed to *ali1-1* homozygotes. In the BC1 F1, 23 recombinant individuals were recovered out of 322 progeny, demonstrating that *ali1* was located 7.1 cM from the *Rp1* locus on chromosome 10S (Table S5). Genomic DNA was sequenced from *ali1-1* mutants derived from nine independent *ali1-1* x B73 BC1 F2 families. Within the mapped region, a single homozygous G to A transition was identified which resulted in a premature stop codon in the last exon of GRMZM2G180205 (v4 gene model Zm00001d023264) removing the last 16 amino acids from the protein-coding sequence (Fig. S7A, S8, S9, and Data table S8). Perfect co-segregation between the mutant phenotype and this G to A transition (*n*=125) was observed in a F2 population derived by crossing *ali1-1* with the maize inbred line M37W. GRZM2G180205 encodes a WD-40 repeat protein and the ortholog of the triple A syndrome (AAAS; OMIM 231550) human disease gene, also called the ALacrimia-Achalasia-aDrenal Insufficiency Neurologic disorder1 (ALADIN1) protein, which is a component of the NPC (Fig. S7B and S9). A large-scale duplication of ∼20kb at this locus resulted in a pseudogene annotated as GRZM2G180249 (v4 gene model Zm00001d023266) and encoding sequences similar to the latter half of the protein sequence of GRZM2G180205. All B73-derived ESTs in NCBI match the sequence of GRZM2G180205 and not GRZM2G180249 at the 10 SNPs and N-terminal deletion that distinguish the paralogs (Table S6). The *ali1-1* mutant (Exon 16, W430*) resulted in an amber stop codon within the final WD-40 domain of ALADIN1. A nonsense allele at the homologous position in the human ALADIN protein encodes a disease allele (Exon 16, W474*) that results in moderate AAAS when inherited in combination with a likely null allele, but has not been observed as a homozygote in human (Houlden et al., 2002).

### ali1 affects plant development and asymmetric cell division in an allelic dosage manner

A targeted EMS mutagenesis approach was used to generate additional mutant *ali1* alleles. Mutants were used as ear parents and crossed with EMS-mutagenized B73 pollen. 10,000 M1 plants were screened for the *ali1-1* phenotype and mutant plants were genotyped at the *ali1-1* allele by dCAPS to distinguish pollen contaminants and gynogenetic haploids (Sarkar & Coe, 1966) from plants harboring novel alleles. Twelve plants displayed phenotypes reminiscent of *ali1-1*, but 11 were homozygous for *ali1-1*. One plant was heterozygous for the *ali1-1* mutation and sequencing of GRMZM2G180205 isolated by PCR from DNA extracted from this plant contained a new C-to-T transition at nucleotide position 2925 in exon 10 resulting in a premature stop codon that would remove the last 134 amino acids (Fig. S7A, S8, and S9). Testcrosses to *ali1-1* and wild-type B73 confirmed that the new allele, *ali1-2*, failed to complement *ali1-1*, was recessive to the wild-type allele, and that *ali1-1/ali1-2* heterozygotes had a phenotype intermediate to the two homozygotes (Fig. 1 F,G). The *ali1-2* homozygotes were severely reduced in height and arrest in development after producing 9-10 leaves, without developing a tassel or ear or undergoing stem elongation (Fig. 1 D, E).

The mutant alleles displayed incomplete dominance to one another, unlike their recessive relationship to the wild-type allele. The *ali1-1*/*ali1-2* heterozygotes were 55% the height of wildtype, 77% the height of *ali1-1* homozygotes (Fig. 1 F,G), and were taller than *ali1-2* homozygotes. In addition, unlike *ali1-*2 homozygotes which fail to produce any seeds, *ali1-1/ali1-2* heterozygotes were fertile. Given the greater severity of the *ali1-2* phenotype, this suggests that the gene product encoded by the *ali1-1* allele was partially functional and dosage sensitive.

Numerous defects in leaf morphology were visible in *ali1-1* and *ali1-2* mutants. Some *ali1-2* mutant leaves split and became necrotic at the midrib as they aged. Both *ali1-2* and *ali1-1* leaf surfaces were crinkly or wrinkled in appearance (Fig. 1). To investigate the basis of this, epidermal peels were taken from one juvenile leaf (leaf 4) and one adult leaf (leaf 10) from greenhouse-grown plants and an adult leaf 8 from field-grown plants. In adult leaves, both *ali1-1* and *ali1-2* mutants had more aberrant stomatal complexes than non-mutant plants, primarily due to aberrant or a lack of subsidiary mother cell (SMC) divisions, and *ali1-2* was more severely affected than *ali1-1* (Fig. 2 A-D). Inheritance of this leaf trait was monogenic recessive (Fig. 2 E-G). There were no discernable effects on nuclear shape in epidermal subsidiary or pavement cells of *ali1-1*/*ali1-1* plants compared to wild type when observed at maturity with 4′, 6-diamidino-2-phenylindole (DAPI) staining (Fig. S10 and S11). Consistent with the greater severity of defects in adult tissues, the *ali1-1* mutants did not discernably alter SMC division in the juvenile fourth leaf. The more severe *ali1-2* mutant, on the other hand, did affect aberrant SMC divisions in juvenile leaves indicating that the *ali1* gene was necessary for normal juvenile stomatal development.

**Fig. 2.**
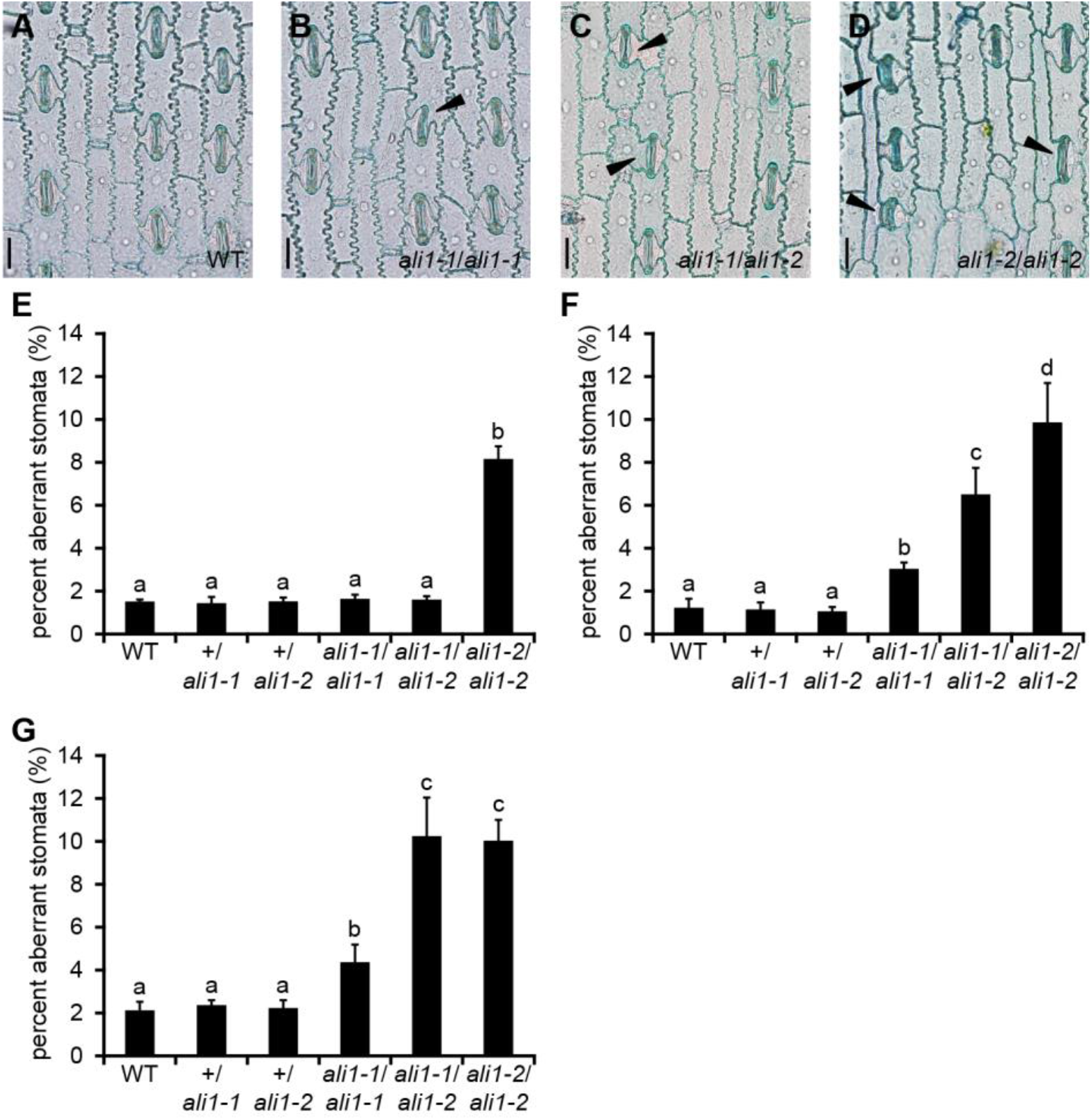
Aberrant stomatal complexes in *ali1* mutants. (**A-D**) Leaf epidermal peels stained with TBO of the 10^th^ leaf of (**A**) wild-type, (**B**) *ali1-1*/*ali1-1*, (**C**) *ali1-1*/*ali1-2*, and (**D**) *ali1-2*/*ali1-2* plants at V11 stage in the greenhouse. Black triangles indicate subsidiary cells of aberrant stomata. Scale bars, 20µm. (**E-G**) The percentage of aberrant stomata in wild type (*n*=6), +/*ali1-1* (*n*=6), +/*ali1-2* (*n*=6), *ali1-1*/*ali1-1* (*n*=6), *ali1-1*/*ali1-2* (*n*=6), and *ali1-2*/*ali1-2* (*n*=6) from the (**E**) 4^th^ leaf at V5 stage or (**F**) 10^th^ leaf at V11 stage grown in the greenhouse or (**G**) from the 8^th^ leaf at V9 stage grown in the field. (**E-G**) Error bars indicated standard deviation. Connecting letter report indicating significance between genotypes determined by ANOVA and post-hoc analysis using the Holm-Sidak algorithm.

Incomplete dominance between the mutant alleles was also observed for the SMC division phenotypes. Normal SMC divisions were observed on the fourth leaf of *ali1-1*/*ali1-2* plants grown in the greenhouse indicating that the protein encoded by *ali1-1* was sufficient to support normal SMC division in juvenile leaves and *ali1-1* was dominant to *ali1-2* (Fig. S12 A-F). A greater proportion of stomatal complexes were aberrant on the eighth leaf of field grown (Fig. S12 G-L) and the tenth leaf of greenhouse grown *ali1-1*/*ali1-2* plants than in *ali1-1* homozygous mutants, indicating that a single dose of *ali1-1* was insufficient to support normal development after phase change. The *ali1-1*/*ali1-2* heterozygotes were still less severe than the *ali1-2* homozygotes in the greenhouse-grown adult tenth leaves indicating the same dosage dependence as observed for height (Figure 1). Interestingly, the SMC division phenotypes of *ali1-1/ali1-2* heterozygotes and *ali1-2* homozygotes were not different on the eighth leaf of field grown plants, indicating that *ali1-2* was completely dominant to *ali1-1* for SMC division in adult leaves in the field but not greenhouse conditions. The *ali1-1* mutant plants are suppressed in the greenhouse and the reduction in aberrant stomata of *ali1-1*/*ali1-2* as compared to *ali1-2* when grown in the greenhouse is likely due to differences between the two growing environments (Fig. 1 and 2). The aberrant SMC divisions were not accompanied by defects in stomatal initiation as there were no significant differences in stomatal indices or stomatal densities (Table S7 and S8). Therefore, the SMC division defect was solely the result of failed asymmetric division and not due to an increased number of guard mother cells or SMCs.

### NPC members show a transcriptional compensation effect in ali1-1 mutants

To investigate the gene expression consequences of reduced *ali1* function we performed RNA-seq experiments comparing *ali1-1* and wild type tissues. Tassels were the most severely affected organ in *ali1-1* mutants (Fig. 1). RNA was extracted from stem tissue in developing tassels just before tassel emergence from the developing leaf whorl from wild-type and mutant plants and differential gene expression was assessed by RNA-seq. Assessment of differential gene expression identified 962 genes significantly up-regulated and 1719 genes down-regulated in *ali1-1* tassels as compared to wild type with FDR less than 0.05 and greater than two-fold change (Data table S2). The ALADIN1 transcript was not differentially expressed in the experiment. The presence of the ALADIN1 transcript in the *ali1-1* mutant provides an explanation for apparent partial function of the *ali1-1* allele and indicates that the premature stop codon that was still encoded by the last exon did not trigger nonsense-mediated decay of the mRNA. Known determinants of asymmetric division in the SMC, PANGLOSS1 (PAN1) and suppressor of cAMP receptor/Wiskott-Aldrich syndrome protein-family verprolin-homologous protein (SCAR/WAVE) complex member BRICK1 (BRK1) (Facette et al., 2015; Gallagher & Smith, 2000), were significantly down-regulated in *ali1-1* tassels (Data table S9) suggesting that mis-expression of these genes may contribute to aberrant control of SMC divisions in *ali1* mutants. Among the differentially expressed genes, only four NPC genes, RIBONUCLEIC ACID EXPORT1-LIKE1 (RAE1-L1), CANDIDATE GENE1 (CG1), NUCLEOPORIN205 (NUP205), and CONSTITUTIVE EXPRESSOR OF PR GENES5 (CPR5) were significantly up-regulated in *ali1-1* tassels (Fig. 3 and Table S9). RAE1 and ALADIN from *A. thaliana* physically interact and CG1 was previously predicted to be in the same subcomplex as ALI1 on the cytoplasmic region of the NPC (Tamura et al., 2010). To assess if other ALADIN-interacting proteins might display compensatory changes in expression in the *ali1-1* mutant, genes similar to known ALADIN1-interacting proteins from humans were identified. Of the 64 maize sequences similar to the human ALADIN protein interactors, six were differentially expressed genes at a false discovery rate (FDR) of <0.05 including the C-TERMINAL DOMAIN NUCLEAR ENVELOPE PHOSPHATASE1-L1 (CTDNEP1-L1), RAN BINDING PROTEIN2-L5 (RANBP2-L5), and RANBP2-L7 genes (Table S10). ALADIN has been shown in humans to be anchored to the NPC via NUCLEAR DIVISION CYCLE1 (NDC1) (Gu et al., 2016; Kind, Koehler, Lorenz, & Huebner, 2009; Yamazumi et al., 2009) but a maize homolog of this gene was not differentially expressed in *ali1-1* mutants. The NPC is comprised of ∼30 subunits that have been described in detail in Arabidopsis (Tamura et al., 2010) so we identified the maize genes encoding the homologs of these genes (Fig. 3, Table S7, and Table S8). In developing mutant tassels, the mRNA encoded by NPC component genes were more likely to be accumulated at higher levels (40 of 49 expressed transcripts) in *ali1-1* mutant tassel stem tissue than in wild type (χ^2^ p-value = 9.49 E-06) (Fig. 3).

**Fig. 3.**
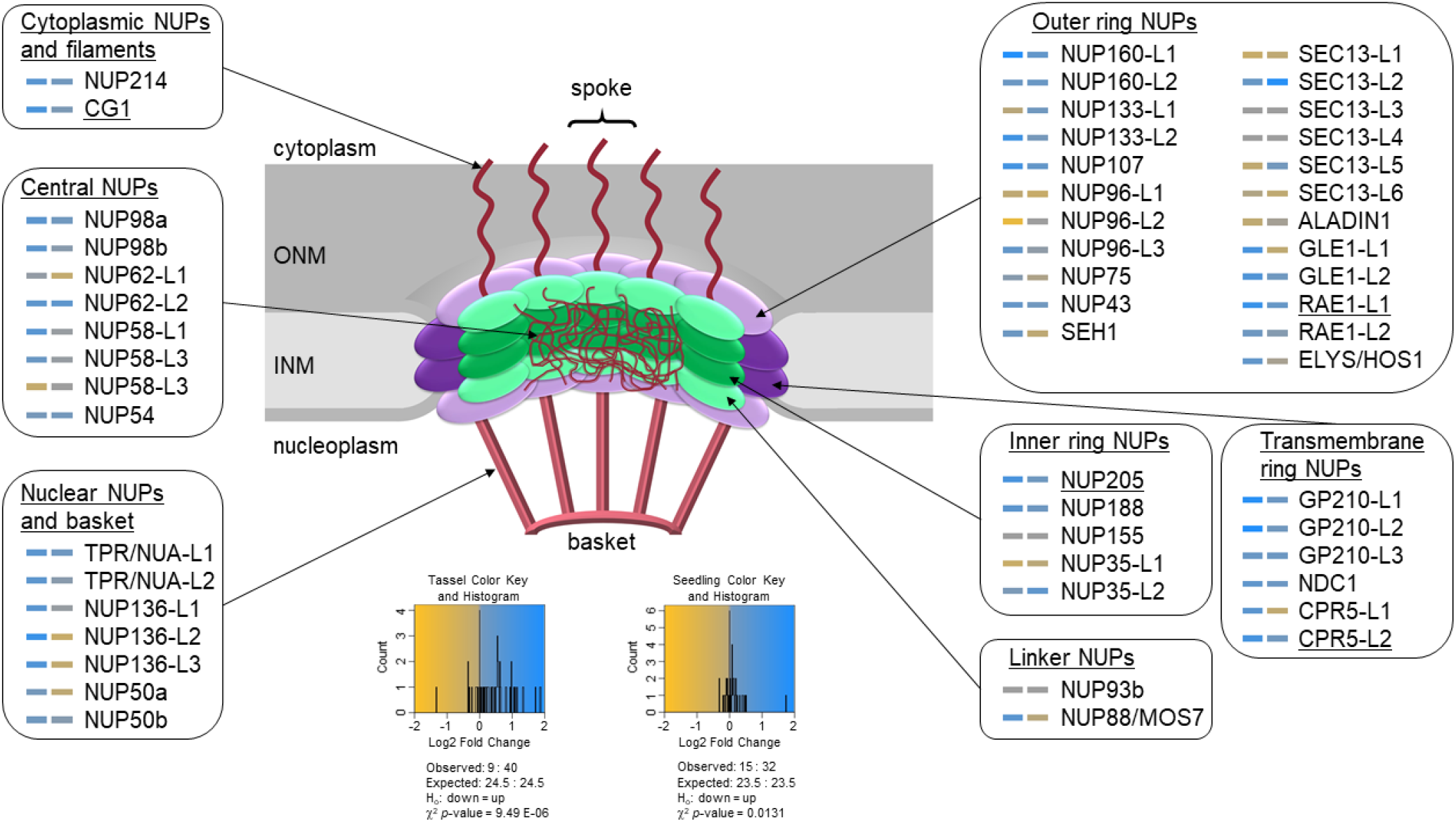
NPC gene expression in *ali1-1* mutants. Heatmap of NPC gene expression in *ali1-1* tassels (left) and *ali1-1* seedlings (right) as compared to respective wild type. Putative NPC genes in maize were identified by BLASTP to NPC members from Arabidopsis (Kind et al. 2009; Tamura et al. 2010; Tamura and Hara-Nishimura 2013; Gu et al. 2016). Outlined gene groups indicate sub-complexes within the NPC as relative location and structure is shown in the middle figure. Gold color indicates down-regulation and blue indicates up-regulation of respective transcript as indicated by the color key. Underlined transcripts (CG1, RAE1-L1, NUP205, and CPR5-L2) were significant at genome-wide Benjamini-Hochberg corrected *P* < 0.05 for *ali1-1* tassels (left). The observed and expected down-regulated and up-regulated transcripts in the NPC, hypothesis test, and χ^2^ *p*-value are shown below the color key and histogram for both the tassel and seedling data.

Unlike tassels, juvenile leaves were the least phenotypically affected aerial tissues in the *ali1* mutants. In an effort to better assess direct effects of *ali1-1*, we performed RNA-seq using RNA extracted from mutant and wild-type leaves at 15 days after planting (DAP), before any phenotypes were visible (Fig. 2 E and Fig. S12 D). Again, more NPC genes were up-regulated than down-regulated in *ali1-1* as compared to wild type (32 of 47 expressed transcripts; χ2 p-value = 0.0131; Fig. 3, Table S11, and Data table S10). However, only 16 genes were significantly up-regulated and 40 genes down-regulated after a transcriptome-wide FDR adjustment and this dropped to 9-up and 14-down when a two-fold expression difference threshold was applied (Data table S3). This indicates that the weak *ali1-1* allele triggered the accumulation of transcripts encoding NPC components prior to visible effects on plant growth or SMC division defects. Of the maize homologs of ALADIN interactors in human, only RANBP2-L4 was differentially expressed (Table S11).

## Discussion

The *ali1* mutants, identify a member of the NPC complex as critical for asymmetric division in plants. A similar effect on subsidiary cell asymmetric division was recently observed for the *mlks2* mutant in maize; which is a component of the LINKER OF NUCLEOSKELETON AND CYTOSKELETON (LINC) protein complex that spans the nuclear envelope but is not associated with the NPC (Gumber et al., 2019). The MLKS2 transcript was not differentially accumulated in *ali1-1* mutants, nor were any other members of the LINC complex, including SUN- and KASH-domain transcripts (Data Table S2). The LINC complex also affects nuclear shape in plants (Meier, Griffis, Groves, & Wagner, 2016; Moser, Kirkpatrick, Groves, & Meier, 2020; Newman-Griffis, Del Cerro, Charpentier, & Meier, 2019; Thorpe & Charpentier, 2017; Zhou, Graumann, Evans, & Meier, 2012; Zhou, Graumann, & Meier, 2015; Zhou, Groves, & Meier, 2015; Zhou & Meier, 2013) and effects on nuclear shape are observed in the *mlks2* mutant (Gumber et al., 2019) but were not observed in *ali1-1* mutant leaves. To date, only one mutant in a nuclear pore component, Nup136 in Arabidopsis, controls the shape and size of nuclei (Tamura & Hara-Nishimura, 2011). In addition to our observations of normal nuclear shape in *ali1-1*, no other studies of ALADIN1 homologs in other species identified an effect on nuclear morphology (Huebner et al., 2009; Huebner et al., 2006).

The aberrant cell divisions in the *ali1* mutants suggest that the NPC can affect nuclear localization or asymmetric phragmoplast assembly at the site of asymmetric divisions. Other mutants of maize, in addition to *mlks2*, result in aberrant SMC divisions. Among these are mutants in genes encoding components of the actin-like protein containing SCAR/WAVE complex (Djakovic, Dyachok, Burke, Frank, & Smith, 2006; Facette et al., 2015; Frank, Cartwright, & Smith, 2003). Interestingly, the SCAR/WAVE component BRK1 mRNA is decreased in *ali1* mutants. The *brick1* mutant results in a loss of epidermal crenulation, which are more evident in adult leaves, and aberrant SMC divisions. Similarly, PAN1 mRNA is decreased in *ali1* (Table). The PAN1 protein is localized via the SCAR/WAVE complex and, mutants of *pan1* are also affected in asymmetric division and produce aberrant SMC. Given the similar impacts on SMC divisions and the gene expression effects, *ali1* may link the SCAR/WAVE complex, required for actin polymerization and cytoskeletal dynamics, to nuclear positioning and asymmetric division control. Exploration of the relationship between ALI1 and cell division dynamics awaits future experimentation in maize as well as in human where these processes may contribute to the etiology of AAAS.

Identification of the weak loss of function *ali1-1* allele and the stronger *ali1-2* allele allowed us to investigate the dosage-dependent effects on phenotypes. Plant height and SMC division displayed variation in phenotypic expression across an allelic series. The *ali1-1* and *ali1-2* alleles were completely recessive when in combination with the wild-type allele, indicating that one wild-type copy was sufficient to maintain NPC function. The *ali1-1*/*ali1-2* heterozygote was more severe, for both plant height and SMC asymmetric cell division, than *ali1-1*/*ali1-1* mutants indicating that *ali1-1* was haploinsufficient with the *ali1-2* allele. These effects were more obvious in adult tissues, adult leaves and stems, and were consistent with an increasing demand for ALI1 after phase change. The observed increase in other phenomena affected by SCAR/WAVE action, such as leaf crenulation, after phase chance may indicate a greater accumulation or role for this protein complex in epidermal development of adult leaves. If this is the case, greater accumulation of ALI1 may be required for stoichiometric protein-protein interactions. This has the capacity to create the threshold effect on SMC divisions at the higher levels of ALI1 activity provided by the *ali1-1* mutant homozygotes. Further investigation into these allelic combinations at the protein level and their interactions with determinants of nuclear positioning and the other members of the NPC are needed to test this hypothesis.

Substantial differences were observed for the weak *ali1-1* mutant phenotype depending on whether they were grown in the field or the greenhouse. In repeated seasons, and plantings, *ali1-1* was substantially suppressed and nearly indistinguishable from wild-type siblings when grown in the greenhouse. Though we do not know the cause of this environmentally-contingent phenotype expression in *ali1-1* it is reasonable to propose light quality and day length as factors. The *dracula2* mutant in Arabidopsis encodes a NUP98 ortholog and is involved in shade-avoidance regulated gene expression (Gallemi et al., 2016). Future investigation into the effects of the shade-avoidance response for *ali1* needs to be done.

In this study, we show that the *ali1* mutant was encoded by the maize ortholog of the human disease gene *AAAS*. The *ali1-1* allele is encoded by mutation of the codon for the tryptophan at position 430 of the maize protein to a stop. Remarkably, the same position in the human protein (W474) encodes a known disease allele that is also the result of a nonsense mutation (Houlden et al., 2002). The human and mouse genes encode a C-terminal extension (Fig. S9) but no function of this region has been identified. Deletions of the C-terminal extension after the homologous *ali1-1* position that remove all (Q490*) or part (R493*; V497*; or R500*) of the C-terminal extension did not alter protein localization; however, a R478* allele which truncates fewer amino acids than the maize *ali1-1* mutation, did result in mis-localized GFP-ALADIN in human cells (Janet M. Cronshaw & Matunis, 2003), and resulted in human disease phenotypes. In humans this allele has only been clinically observed in heterozygous condition and results in relatively mild disease in combination with a more severe mutant allele. Given the incomplete dominance observed between *ali1-1* and *ali1-2* mutants, this suggests that this allele is also haploinsufficient in humans (Houlden et al., 2002). It is unclear if homozygotes at the human W474* allele would exhibit weaker, perhaps even subclinical, phenotypes consistent with AAAS. The dramatic phenotypic impacts, and allelic series, available in maize mirror the situation in humans. Further investigation into the transcriptional and cell biological consequences of *ali1-1* and *ali1-2* may help identification of the cellular mechanisms responsible for the progressive deleterious effects of mutating *ali1* and provide a model system for understanding AAAS in human.

Previously identified Arabidopsis mutants defective in NPC subunits alter nuclear transport (Parry, 2014). NPC members also influence gene expression via binding chromatin, changing the chromatin localization in the nucleoplasm, affecting nuclear export of mRNA, and shuttling of proteins (Parry, 2015). The increased accumulation of mRNA encoding NPC subunits in *ali1-1* mutants suggests the existence of an NPC surveillance mechanism that can increase the abundance of multiple transcripts encoding NPC subunits when the complex was compromised. This effect was visible in *ali1-1* mutants at the juvenile stage prior to any visible phenotypic effects suggesting that it was not a secondary response to altered cellular morphology in the mutants. In addition, the lack of a phenotype in juvenile plants may be the result of this compensatory change in NPC subunit expression. If so, a mutagenesis experiment should recover enhancers of the *ali1-1* phenotype when this process is disrupted. It remains to be tested if the observed compensation in NPC subunit transcripts in *ali1-1* are the result of monitoring for aberrant transport processes, as nuclear shape was unaltered in *ali1-1* (Fig. S10 and S11).

## Acknowledgments and comments

Seeds of *ali1-1* and *ali1-2* are available at the Maize Genetics COOP. Raw sequence data is available at the Short Read Archive under BioSample accession numbers: SAMN06764664, SAMN06764861, SAMN06765376, and SAMN07187450.

This work was supported by funds from the National Science Foundation to BPD and GJ (PGRP # 1444503), B.S. (CAREER #1054918), and CW (PGRP #1025976). Work by NBB supported by USDA-NIFA fellowships to N.B.B (NIFA #2017-67011-26077; #2019-67012-29655), from the U.S. Department of Agriculture, National Institute of Food and Agriculture.

Mention of trade names or commercial products in this publication is solely for the purpose of providing specific information and does not imply recommendation or endorsement by the U.S. Department of Agriculture. The U.S. Department of Agriculture is an equal opportunity provider and employer

We would like to thank Jim Beaty and the crew at the Purdue University ACRE for help with field grown maize used in these studies. This paper is presented as a gift on the occasion of Jim Beaty’s retirement from Purdue in thanks for his support of graduate student research. We would also like to thank Rob Eddy and the crew at the Purdue University Horticulture Growth Facilities with assistance for greenhouse grown maize used in these studies.

## Supplementary Materials

Figures S1-S12

Tables S1-S12

External Databases S1-S10

## Supplementary Materials for

### Other Supplementary Materials for this manuscript includes the following

Data table S1 to S10 as zipped archives:

B73_Mo17_SNPs_CDS.zip

DESEQ2_ALI_TASSEL.zip

DESEQ2_ALI_15DAP.zip

ALI1_REP1_RNA_CALLSNPs.zip

ALI1_REP2_RNA_CALLSNPs.zip

WT_REP1_RNA_CALLSNPs.zip

WT_REP2_RNA_CALLSNPs.zip

ALI1_DNA_CALLSNPs.zip

INTEREST_TASSEL.zip INTEREST_15DAP.zip

**Fig. S1.**
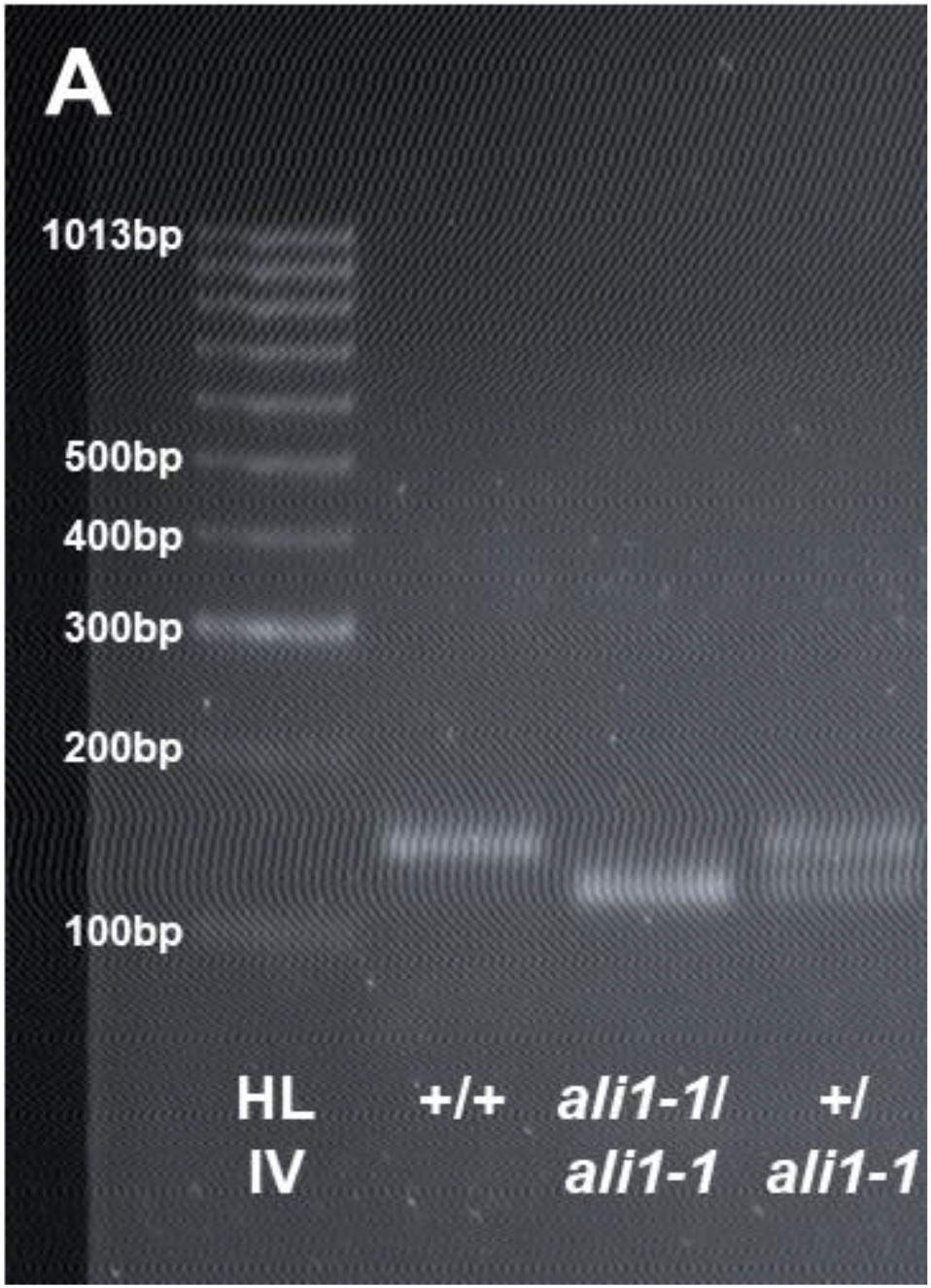
Gel image of genotyping of *ali1-1* allele. After restriction digest of the PCR product using TaqI the wild-type and *ali1-1* mutant bands could clearly be distinguished when run at 95 volts for two hours on a 3% agarose gel. A hyperladder IV standard is shown for size reference (left) with wild-type (left center), *ali1-1* homozygote (right center), and heterozygous (right) PCR product with TaqI digest is shown.

**Fig. S2.**
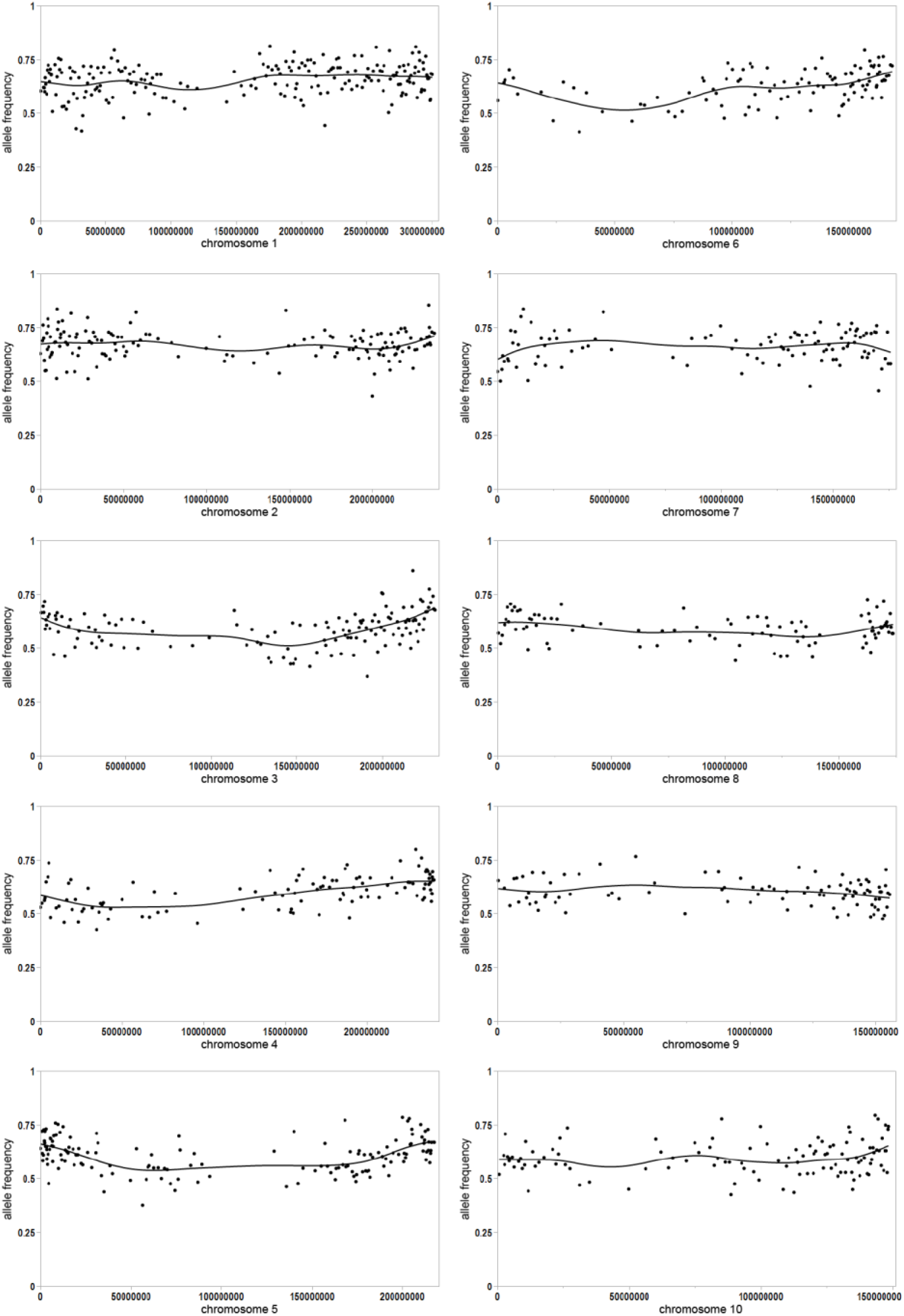
Allele frequency for replicate one of wild-type pools of F2 plants of *ali1-1*/*ali1-1* crossed with Mo17. Allele frequencies of non-overlapping bins of 100 coding sequence SNPs between B73 reference (value of “1” on y-axis) and Mo17 (value of “0” on y-axis) for all 10 maize chromosomes. A cubic spline curve with lambda 0.05 is shown on each plot.

**Fig. S3.**
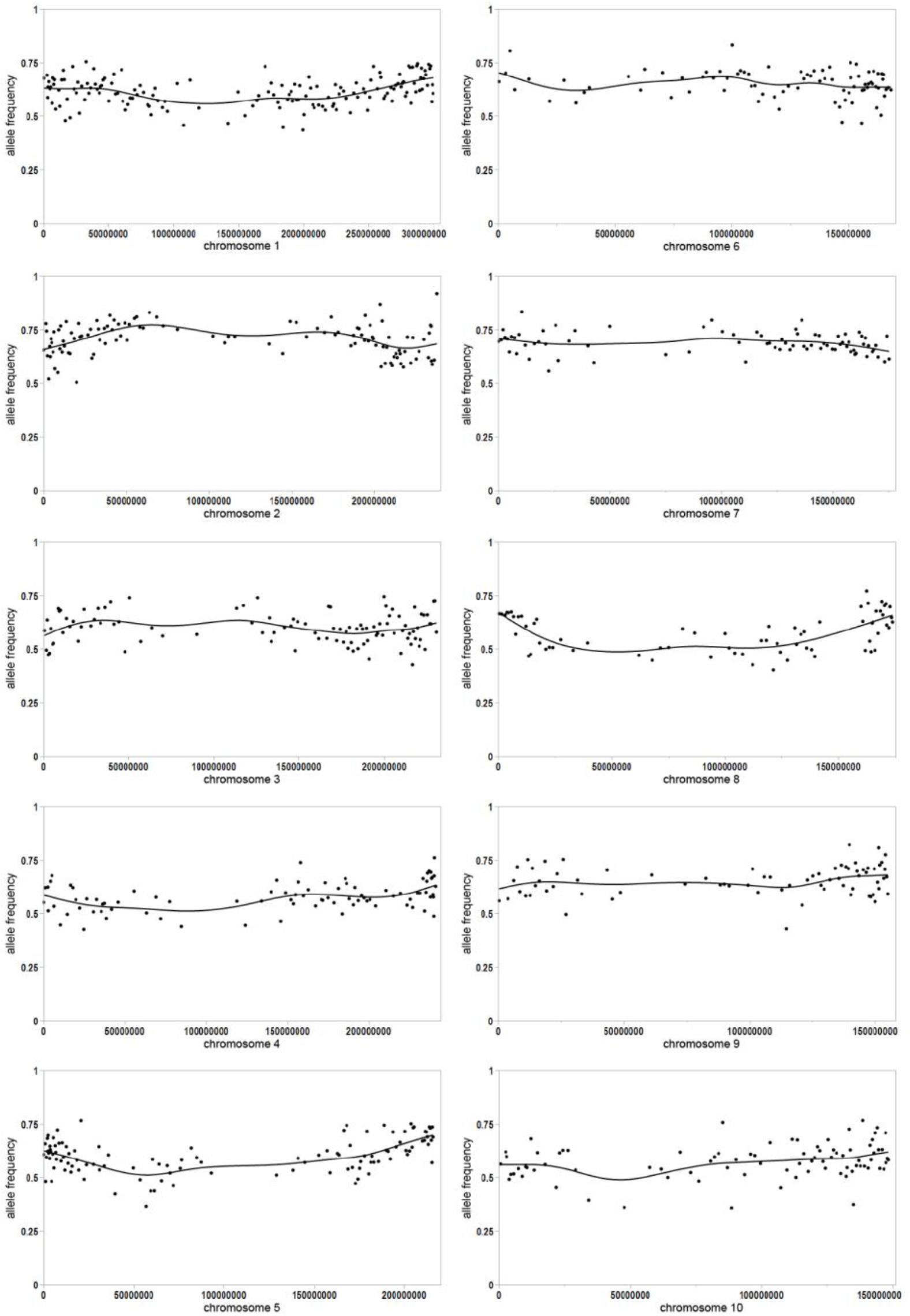
Allele frequency for replicate two of wild-type pools of F2 plants of *ali1-1*/*ali1-1* crossed with Mo17. Allele frequencies of non-overlapping bins of 100 coding sequence SNPs between B73 reference (value of “1” on y-axis) and Mo17 (value of “0” on y-axis) for all 10 maize chromosomes. A cubic spline curve with lambda 0.05 is shown on each plot.

**Fig. S4.**
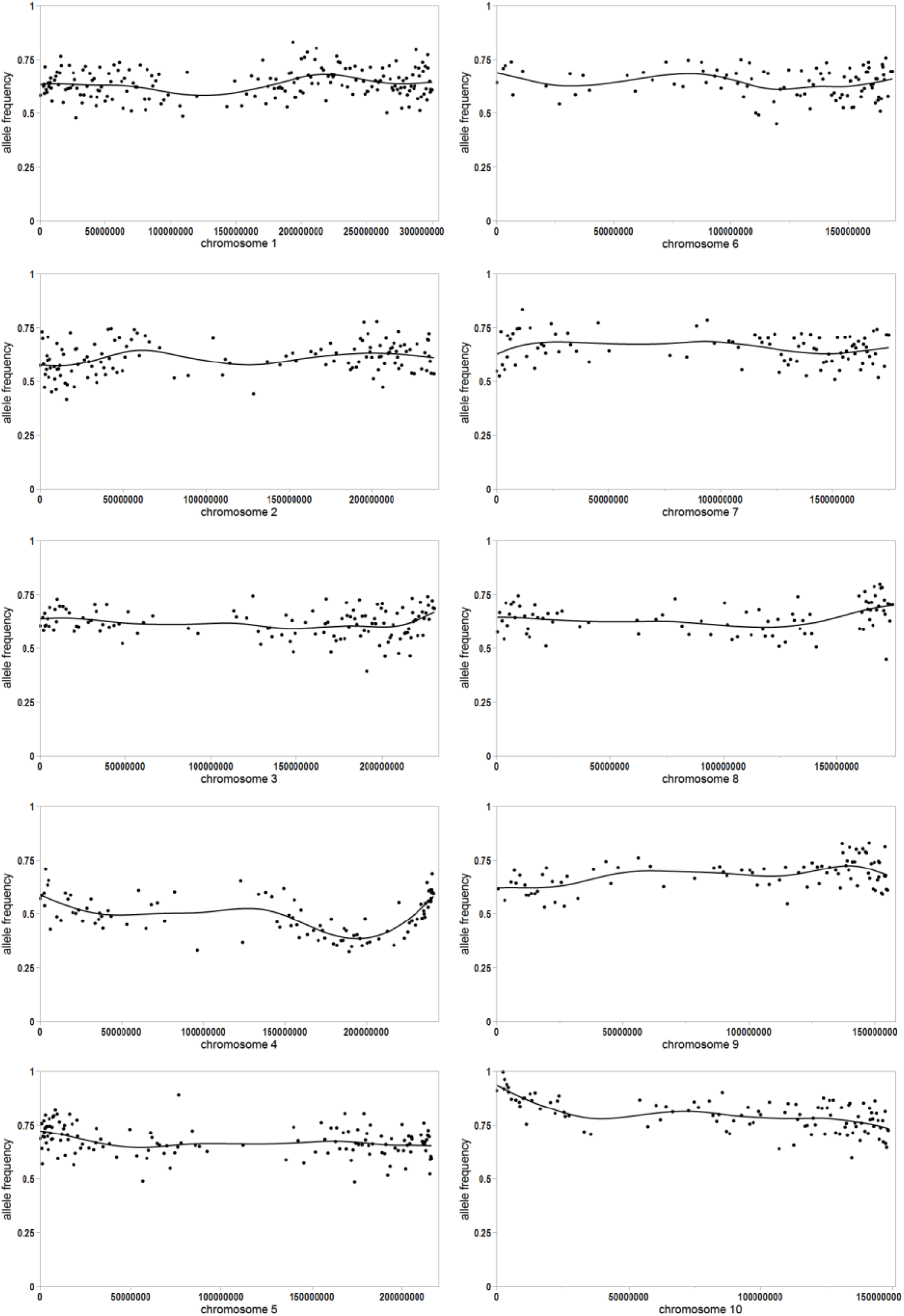
Allele frequency for replicate one of *ali1-1* pools of F2 plants of *ali1-1*/*ali1-1* crossed with Mo17. Allele frequencies of non-overlapping bins of 100 coding sequence SNPs between B73 reference (value of “1” on y-axis) and Mo17 (value of “0” on y-axis) for all 10 maize chromosomes. A cubic spline curve with lambda 0.05 is shown on each plot.

**Fig. S5.**
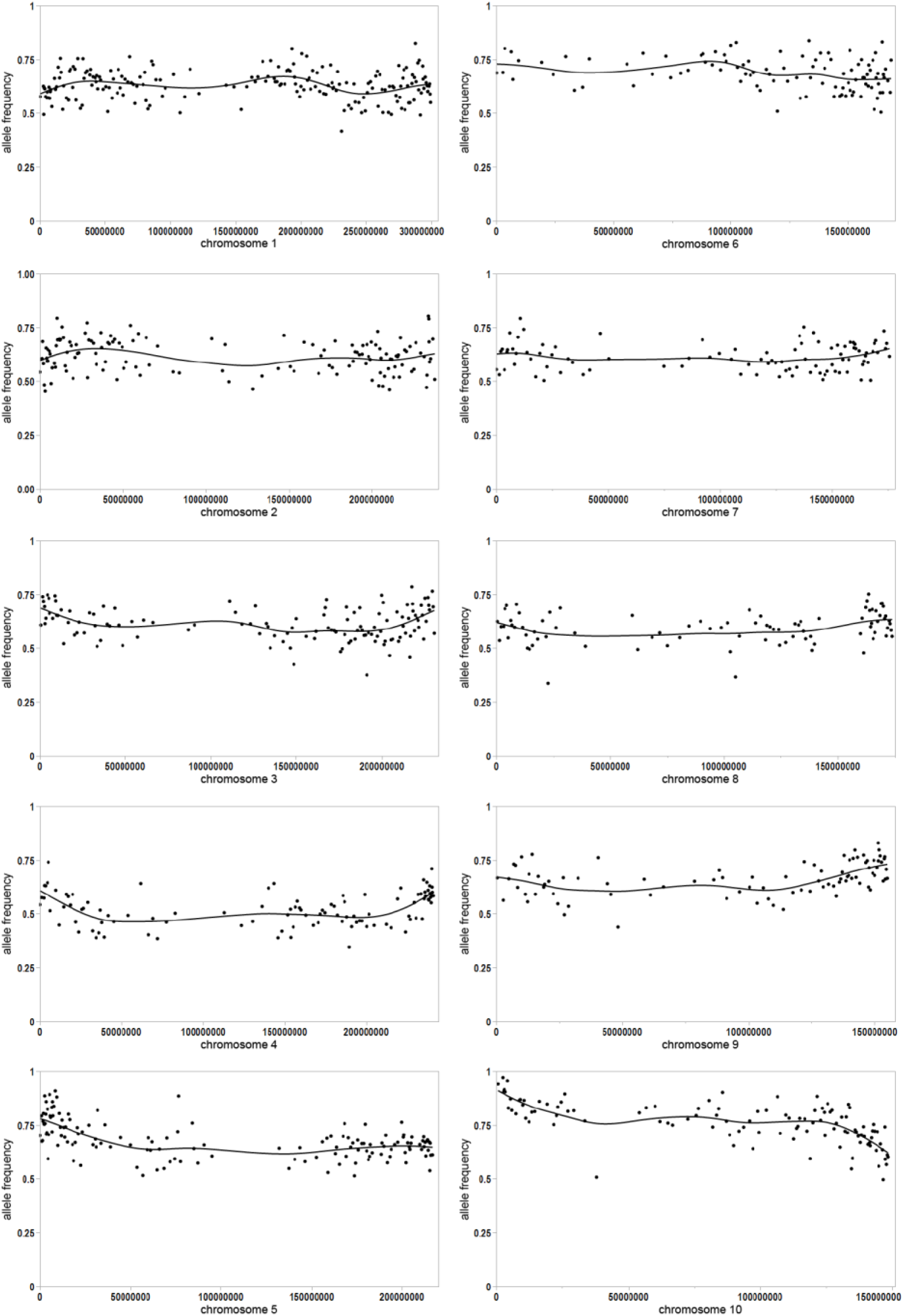
Allele frequency for replicate two of *ali1-1* pools of F2 plants of *ali1-1*/*ali1-1* crossed with Mo17. Allele frequencies of non-overlapping bins of 100 coding sequence SNPs between B73 reference (value of “1” on y-axis) and Mo17 (value of “0” on y-axis) for all 10 maize chromosomes. A cubic spline curve with lambda 0.05 is shown on each plot.

**Fig. S6.**
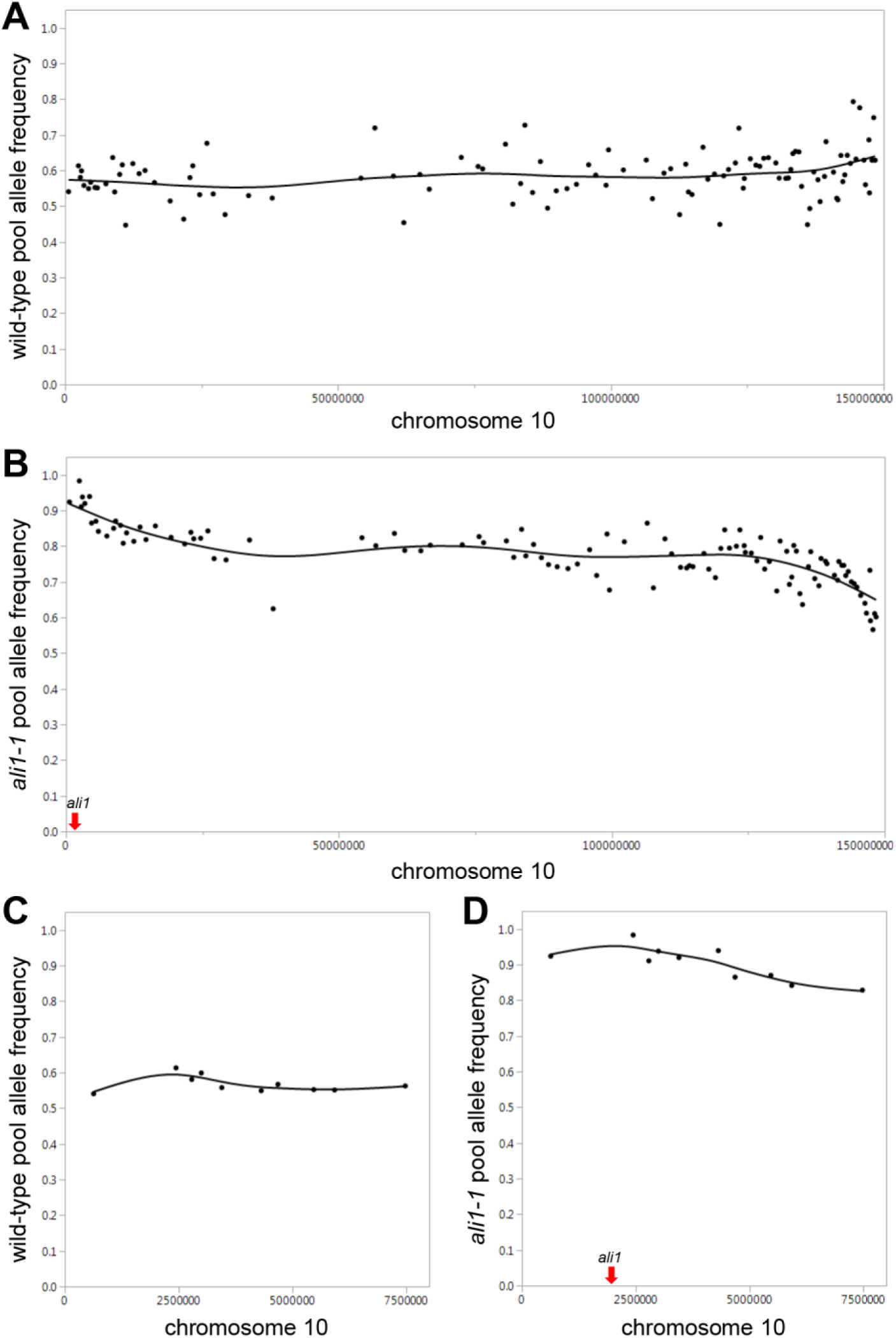
Mapping of the *ali1* gene. SNP allele frequency averages on chromosome 10 from two replicates of pooled (**A**) phenotypical wild-type siblings and (**B**) *ali1-1*/*ali1-1*. SNP allele frequency of *ali1* mapped region from (**C**) phenotypical wild-type sibling pools and (**D**) *ali1-1*/*ali1-1* pools. Allele frequencies are plotted on the y-axis for pools of F2 progeny from an *ali11*/*ali1-1*;B73 x Mo17 self. The y-axis corresponds to the reference genotype frequency in non-overlapping bins of 100 coding sequence SNP per point. The x-axis indicates physical position on chromosome 10. A cubic spline curve with lambda 0.05 is shown. (**B** and **D**) The *ali1* physical position is marked by a red arrow.

**Fig. S7.**
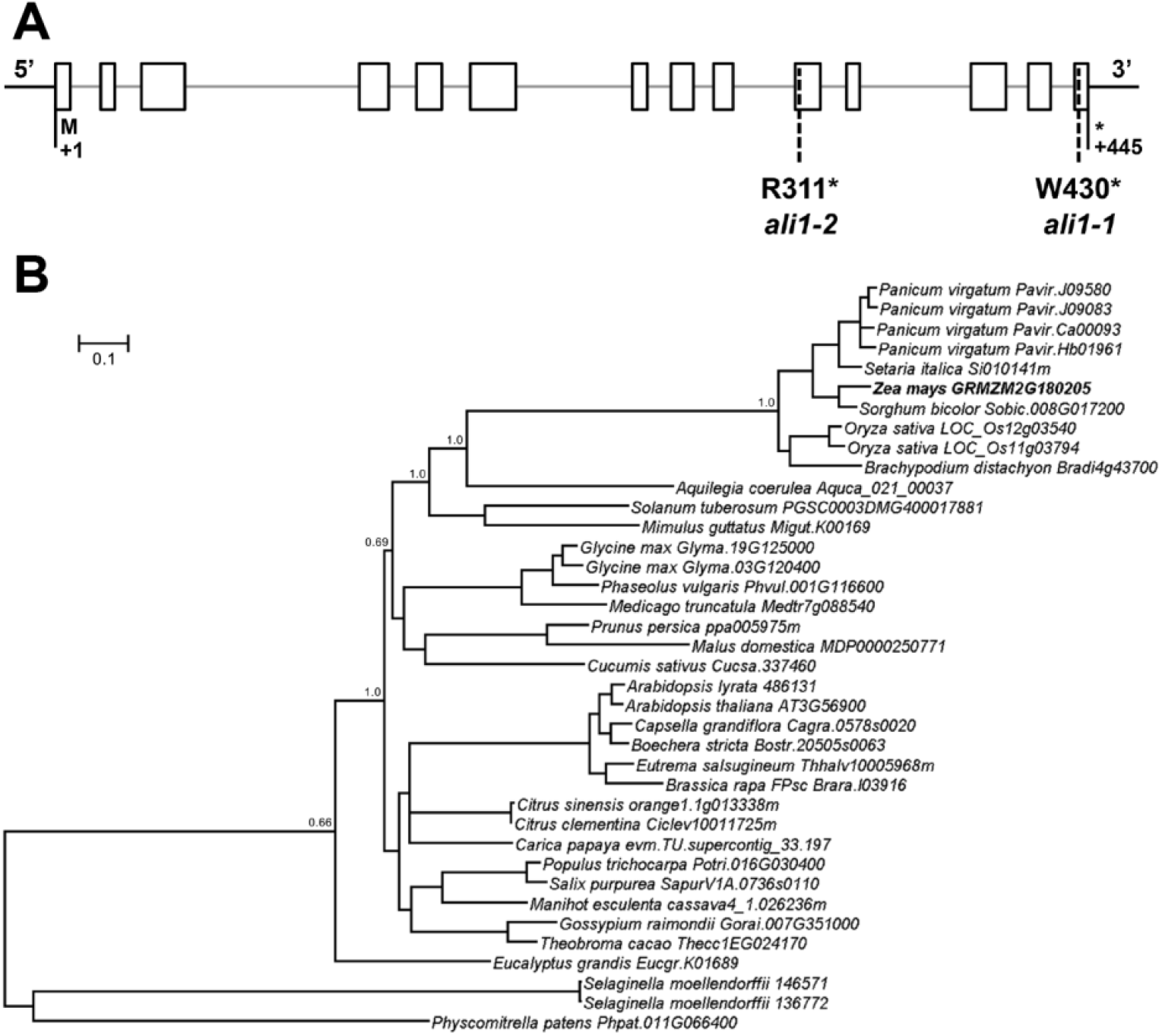
Gene structure of the maize *ali1* gene and phylogenetic tree of *ali1* genes from 32 plant taxa. (**A**) Gene structure of *ali1* and the position of mutations affecting the *ali1-1* and *ali1-2* alleles are shown as dashed lines. Both alleles result in a premature termination codon, as indicated above the allele designations. Exons are represented by white boxes and the introns as grey lines. (**B**) A consensus maximum likelihood, Bayesian Markov chain Monte Carlo, phylogenetic tree of nucleotide coding sequences derived from the *ali1* genes identified within 32 plant genome assemblies in Phytozome. The scale bar indicates the number of codon changes per orthologous site. Posterior probabilities for selected nodes are indicated. The lycophyte species, *Selaginella moellendorffii*, and bryophyte species, *Physcomitrella patens*, were used as an out-group. The maize *ali1* gene is indicated by bold text.

**Fig. S8.**
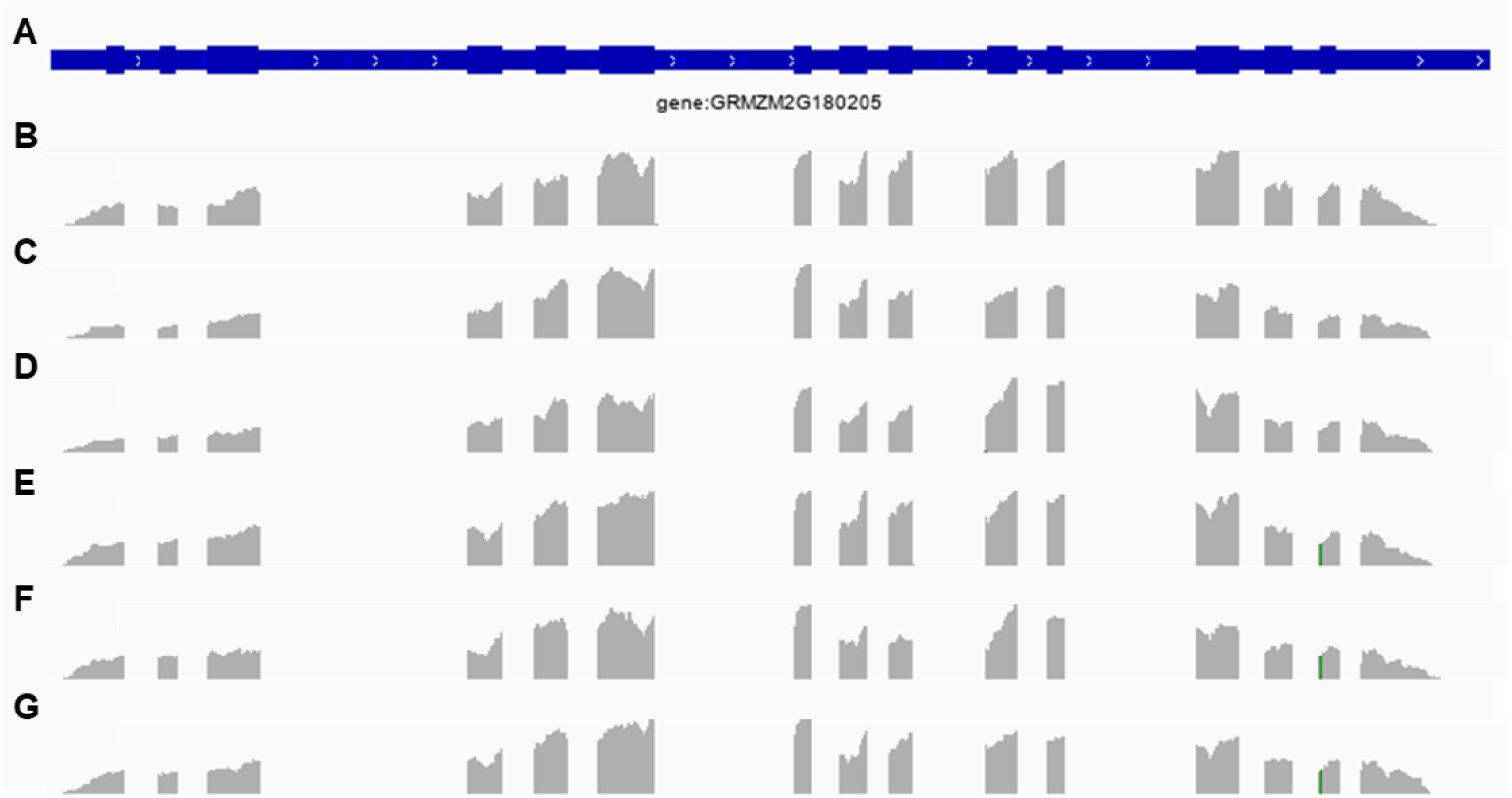
Gene structure of (**A**) *ali1* locus (GRMZM2G180205). Thick portions indicate exons. IGV tracks for RNA reads of WT 15d SAM and surrounding tissue samples (**B**) one, (**C**) two, and (**D**) three. IGV tracks for RNA reads of *ali1-1* SAM samples for replicates (**E**) one, (**F**) two, and (**G**) three. Gray peaks indicate RNA reads aligned to B73 reference genome. (**E**-**G**) Green peak in the 14^th^ exon of *ali1-1* samples indicates causative mutation.

**Fig. S9.**
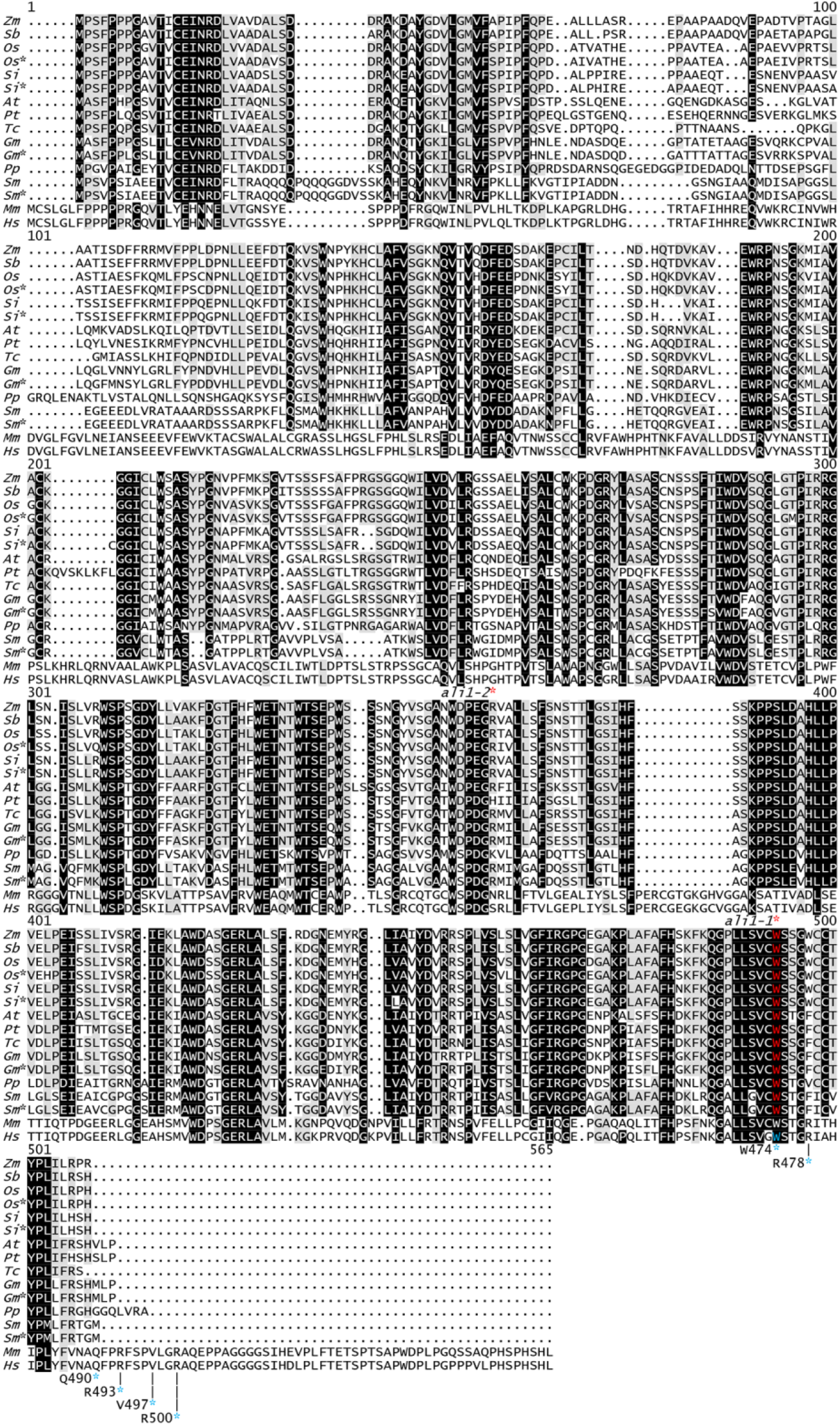
Multiple sequence alignment of the maize *ali1* protein sequence with orthologous sequences. Black background indicates identity conservation of ≥85%, grey specifies conservation of ≥50%, and white background indicates ≤50% conservation. The location of the termination codon in *ali1-1* and *ali1-2* are shown by a red asterisk above the multiple sequence alignment, with the tryptophan codon for *ali1-1* being highlighted in red. The location of previously studied nonsense mutations in *Homo sapiens* and their effects on localization of the ALADIN1 protein are indicated by a blue asterisk below the multiple sequence alignment. Periods designate gaps in the alignment. *Zm, Zea mays* (GRMZM2G180205); *Sb, Sorghum bicolor* (Sobic. 008G017200); Oz, *Oryza sativa* (LOC_Os12g03540); *Oz*, Oryza sativa* (LOC_Os11g03794); Si, *Setaria italica* (Seita.7G310400); *Si*, Setaria italica* (Seita.8G003800); *At, Arabidopsis thaliana* (AT3G56900); *Pt, Populus trichocarpa* (Potri.016G030400); *Tc, Theobroma cacao* (Thecc1EG024170); *Gm, Glycine max* (Glyma.19G125000); *Gm*, Glycine max* (Glyma.03G120400); *Pp, Physcomitrella patens* (Pp3c11_18220); *Sm, Selaginella moellendorfii* (146571); *Sm*, Selaginella moellendorfii* (136772); *Mm, Mus musculus* (uc007xvl.2); *Hs, Homo sapiens* (uc001scr.5).

**Fig. S10.**
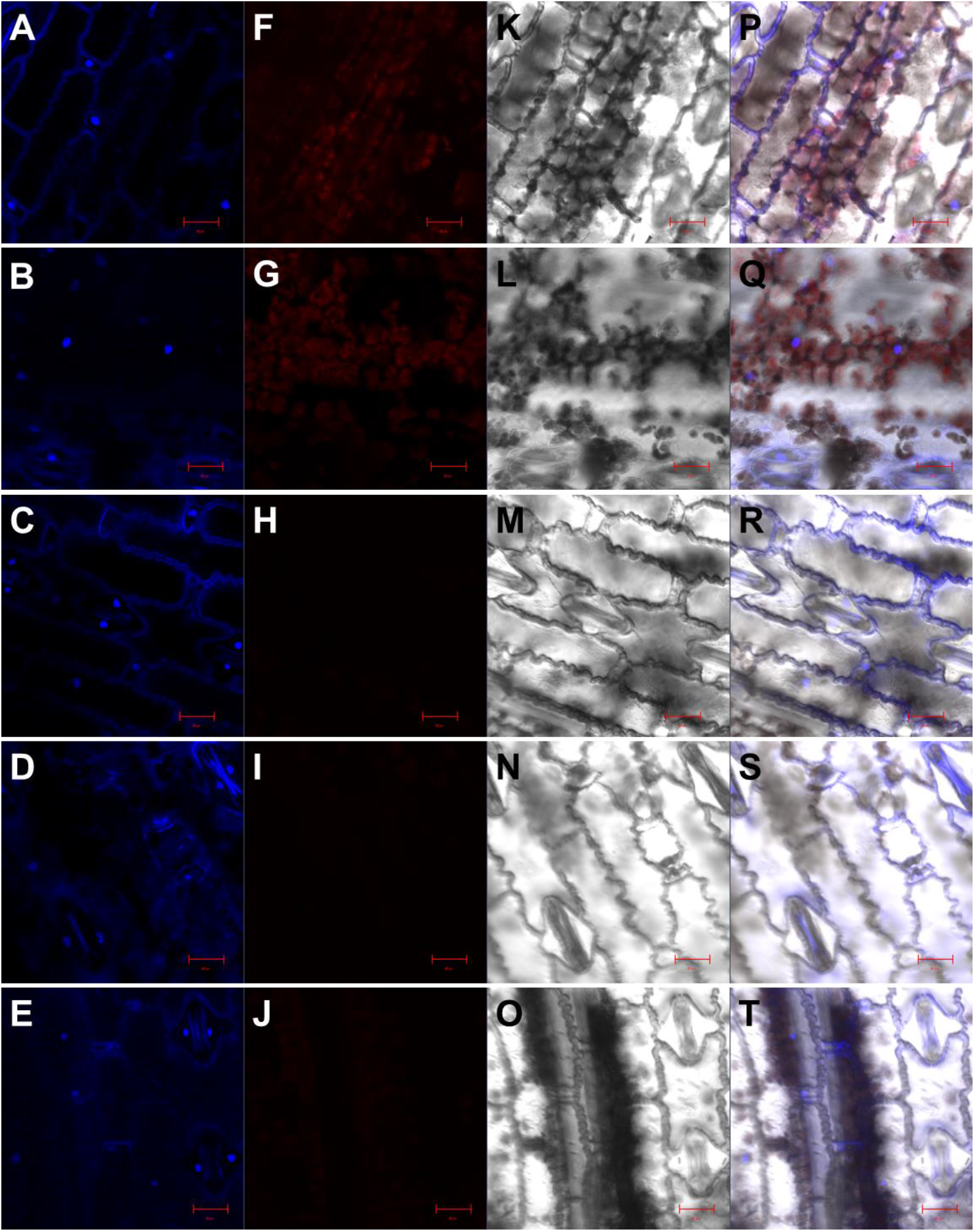
Microscopy of wild-type epidermal cells with 4′, 6-diamidino-2-phenylindole (DAPI) staining to show nuclear morphology. Five replications of wild-type epidermal peels stained with DAPI solution (1 µg/1ml in 1xPBS, pH 8.0) for overnight in dark. DAPI stained epidermal cells were imaged with a Zeiss 880 upright laser scanning confocal microscope. DAPI was excited at 358 nm and visualized with a 461 nm band pass emission filter channel (A-E). Images from each individual sample were also observed with a chlorophyll-a channel (F-J), photomultiplier tube (PMT) channel (K-O), and all three channels merged (P-T). Red scale bars represent 20 µm (A-T).

**Fig. S11.**
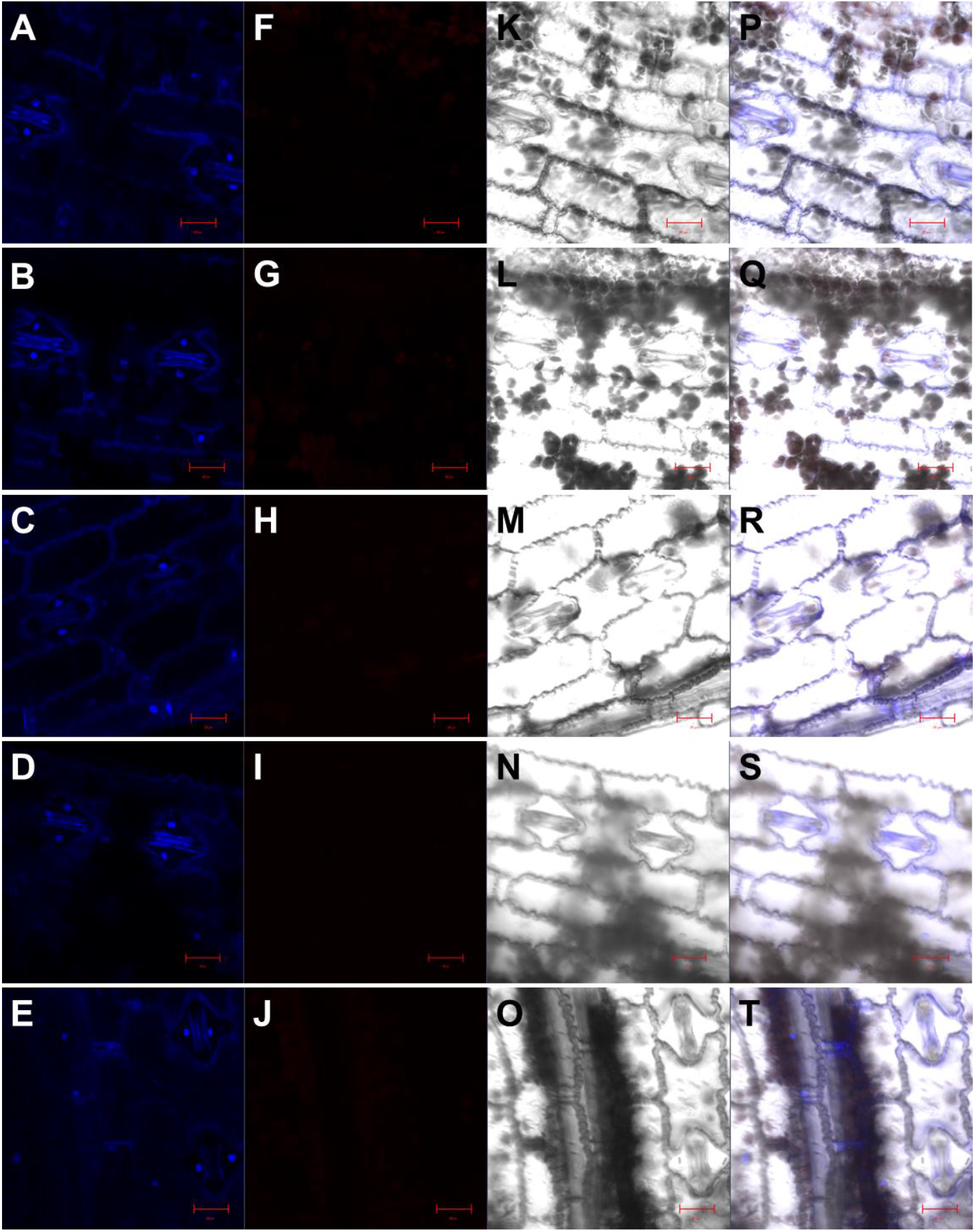
Microscopy of *ali1-1*/*ali1-1* epidermal cells with 4′, 6-diamidino-2-phenylindole (DAPI) staining to show nuclear morphology. Five replications of *ali1-1*/*ali1-1* epidermal peels stained with DAPI solution (1 µg/1ml in 1xPBS, pH 8.0) for overnight in dark. DAPI was excited at 358 nm and visualized with a 461 nm band pass emission filter channel (A-E). Images from each individual sample were also observed with a chlorophyll-a channel (F-J), photomultiplier tube (PMT) channel (K-O), and all three channels merged (P-T). Scale bar represents 20 µm (A-T).

**Fig. S12.**
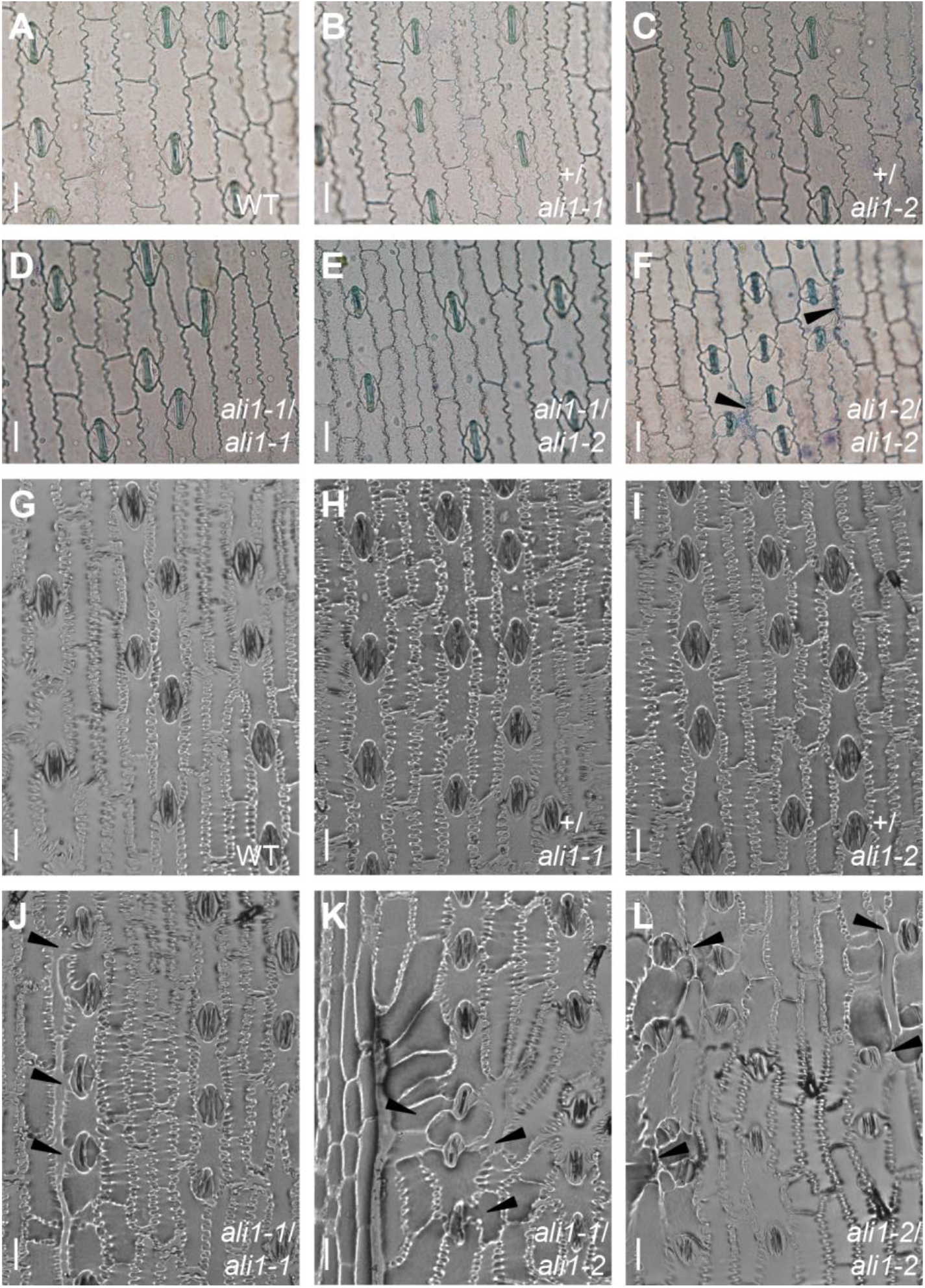
Aberrant stomatal complexes and abnormal cell morphology phenotypes of *ali1* mutants. (A-F) Epidermal peels from the 4th leaf abaxial surface at V5 stage of (A) WT, (B) +/*ali1-1*, (C) +/*ali1-2*, (D) *ali1-1*/*ali1-1*, (E) *ali1-1*/*ali1-2*, and (F) *ali1-2*/*ali1-2* plants grown in the greenhouse. (G-L) Epidermal imprints from the 8th leaf abaxial surface at V9 stage of (G) WT, (H) +/*ali1-1*, (I) +/*ali1-2*, (J) *ali1-1*/*ali1-1*, (K) *ali1-1*/*ali1-2*, and (L) *ali1-2*/*ali1-2* plants grown in the field. (A-L) Black triangles indicate subsidiary cells of aberrant stomata. Scale bar, 20µm.

**Table S1.**
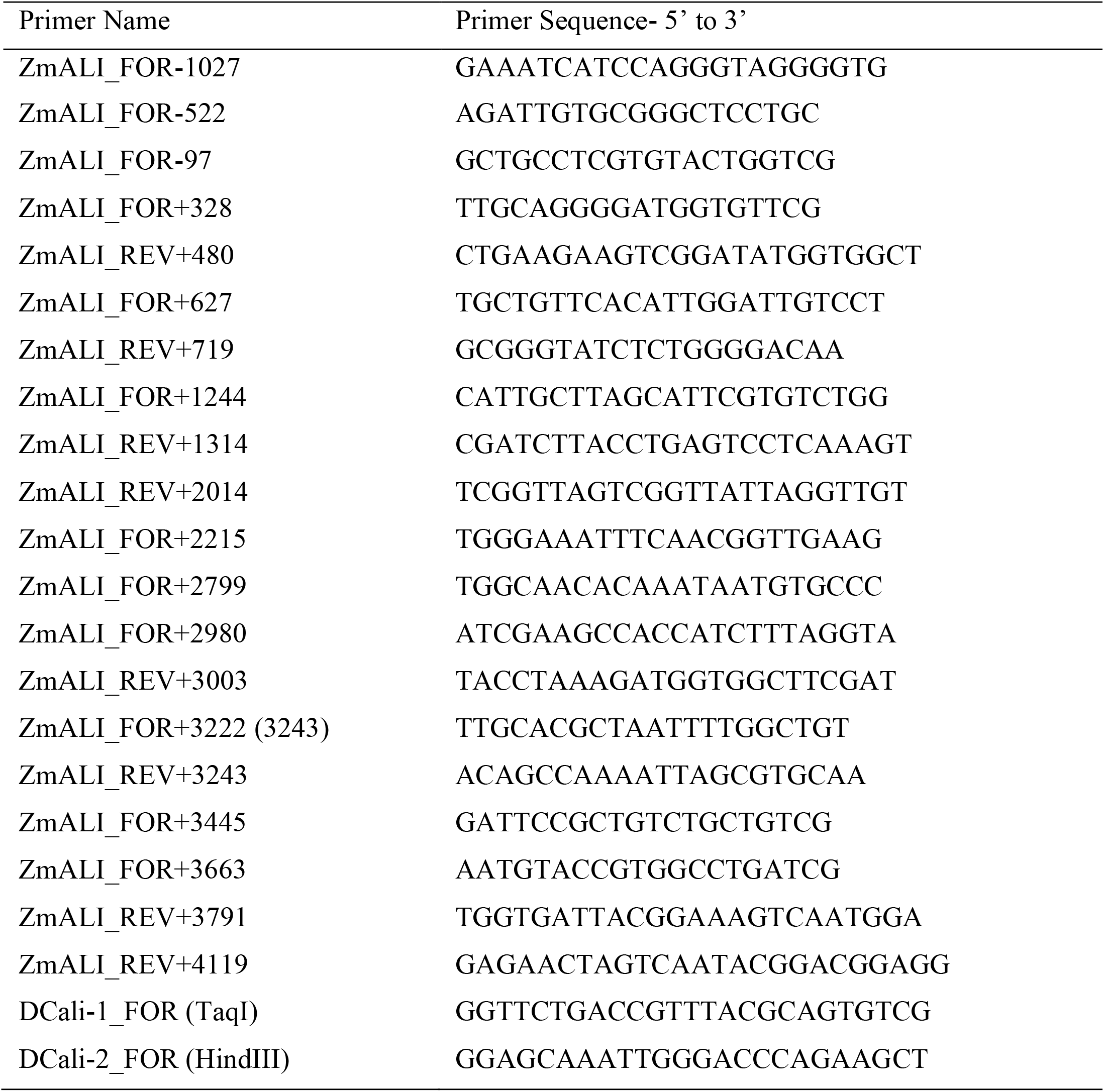
Primers used in this study.

**Table S2.**
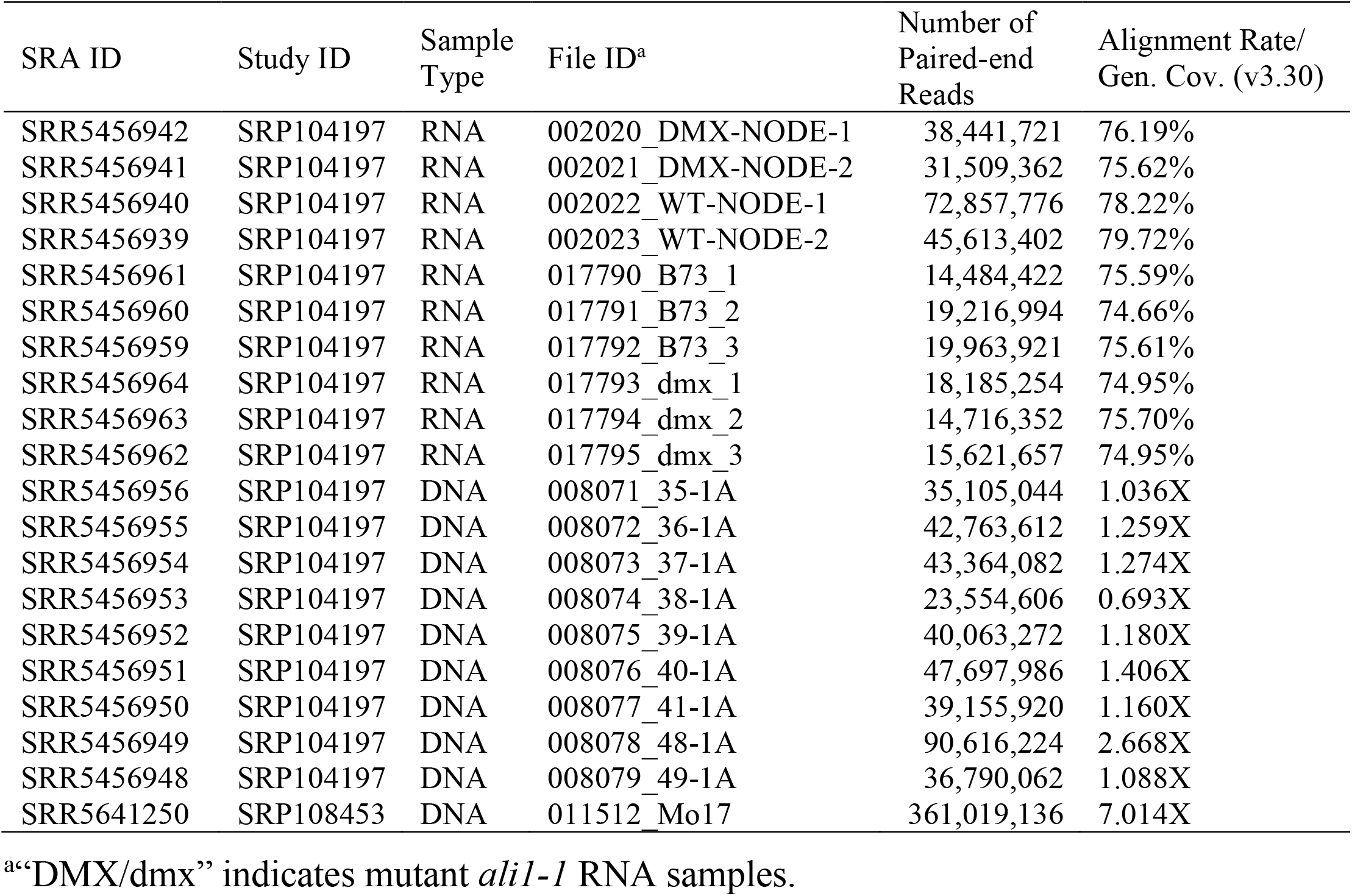
High through-put sequencing information and statistics.

**Table S3.**
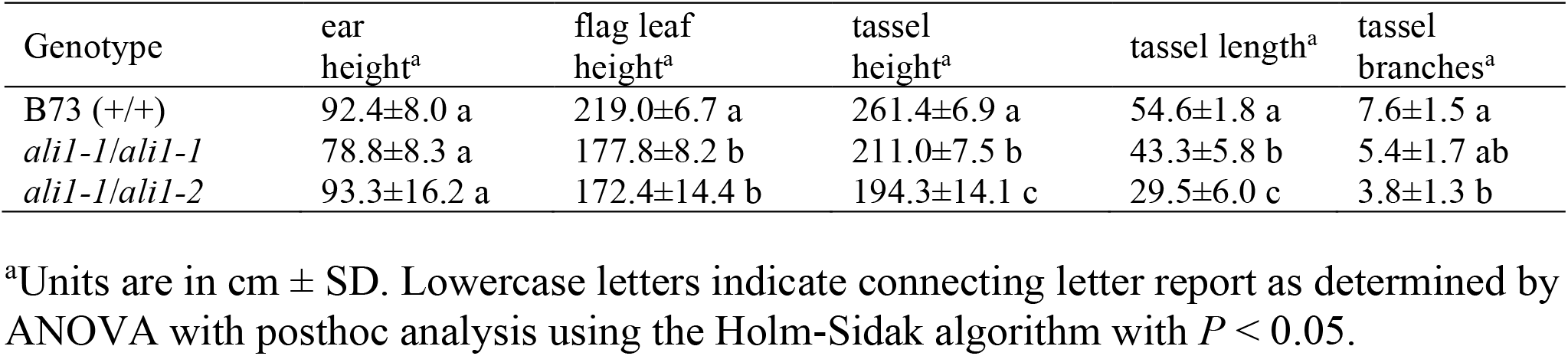
Morphometric analysis of B73, *ali1-1*/*ali1-1*, and *ali1-1*/*ali1-2* grown in the spring of 2016 in greenhouse.

**Table S4.** Plant height of *ali1-1* was suppressed in the greenhouse when grown in the winter of 2015 as compared to field-grown plants.

| | *n* | flag leaf height ± SD<sup>a</sup> |
| --- | --- | --- |
| B73 in greenhouse<sup>b</sup> | 5 | 240.8 ± 12.28 a |
| B73 in field<sup>c</sup> | 16 | 239.5 ± 6.54 a |
| *ali1-1/ali1-1* in greenhouse<sup>b</sup> | 18 | 234.4 ± 14.02 a |
| *ali1-1/ali1-1* in field<sup>c</sup> | 46 | 157.0 ± 19.03 b |

<sup>a</sup>Units are in cm, lowercase letters indicate connecting letter report as determined by ANOVA with post hoc analysis using the Holm-Sidak algorithm with $p < 0.05$ .
<sup>b</sup>Plants were grown in greenhouse in winter of 2015.
<sup>c</sup>Plants were grown in field in summer of 2015 in WL.

**Table S5.** Phenotypic segregation ratios of the progeny of (+/*ali1-1*; +/*Rp1-D21*) crossed with (*ali1-1*/*ali1-1*; +/+) showing linkage disequilibrium between loci.

| Ear | Wild Type | *ali1-1* | *Rp1-D21* | *ali1-1*;<br>*Rp1-D21* | Total | $\chi^2$ *P*-value |
| --- | --- | --- | --- | --- | --- | --- |
| WL15_81 | 0 | 8 | 8 | 1 | 17 | 0.0039 |
| WL15_82 | 1 | 6 | 8 | 1 | 16 | 0.0233 |
| WL15_83 | 0 | 3 | 10 | 0 | 13 | 0.0001 |
| WL15_84 | 0 | 3 | 6 | 1 | 10 | 0.0384 |
| WL15_85 | 0 | 4 | 8 | 0 | 12 | 0.0021 |
| WL15_87 | 2 | 10 | 6 | 1 | 19 | 0.0136 |
| WL15_89 | 1 | 6 | 4 | 1 | 12 | 0.1116 |
| WL15_90 | 0 | 9 | 6 | 1 | 16 | 0.0037 |
| WL15_91 | 0 | 10 | 7 | 1 | 18 | 0.0016 |
| WL15_92 | 0 | 4 | 12 | 0 | 16 | 2.5E-05 |
| WL15_93 | 1 | 7 | 7 | 0 | 15 | 0.0098 |
| WL15_94 | 0 | 5 | 8 | 1 | 14 | 0.0084 |
| WL15_95 | 2 | 6 | 6 | 0 | 14 | 0.0523 |
| WL15_96 | 0 | 7 | 8 | 1 | 16 | 0.0059 |
| WL15_97 | 0 | 7 | 9 | 0 | 16 | 0.0009 |
| WL15_99 | 0 | 8 | 7 | 1 | 16 | 0.0059 |
| WL15_100 | 2 | 5 | 6 | 1 | 14 | 0.1826 |
| WL15_101 | 2 | 9 | 3 | 1 | 15 | 0.0159 |
| WL15_102 | 0 | 7 | 8 | 0 | 15 | 0.0017 |
| WL15_103 | 0 | 9 | 6 | 0 | 15 | 0.0010 |
| WL15_104 | 0 | 7 | 9 | 0 | 16 | 0.0009 |
| WL15_107 | 0 | 4 | 3 | 0 | 7 | 0.0633 |
| Total | 11 | 144 | 155 | 12 | 322 | 3.6E-51 |

**Table S6.** BLASTn results of GRMZM2G180205 genomic sequence against NCBI expression tag sequence database

| Accession Number | Expect value | Percent identity |
| --- | --- | --- |
| DR828860.1 | 2.00E-171 | 100.00% |
| DV511867.1 | 7.00E-122 | 100.00% |
| DV543542.1 | 3.00E-105 | 97.40% |
| FL135742.1 | 2.00E-102 | 99.53% |
| DV029942.1 | 4.00E-99 | 100.00% |
| FL135740.1 | 2.00E-97 | 97.63% |
| DR964374.1 | 7.00E-97 | 100.00% |
| DR828976.1 | 1.00E-93 | 100.00% |
| DR817074.1 | 1.00E-93 | 100.00% |
| EC905516.2 | 9.00E-91 | 98.48% |
| EB407248.1 | 9.00E-91 | 100.00% |
| EE156325.2 | 3.00E-90 | 99.47% |
| EB641592.1 | 3.00E-90 | 100.00% |
| EB165604.1 | 3.00E-90 | 99.47% |
| DV523800.1 | 3.00E-90 | 99.47% |
| DR969589.1 | 3.00E-90 | 99.47% |
| CO468963.1 | 3.00E-90 | 99.47% |
| DN215222.1 | 3.00E-90 | 99.47% |
| CF244328.1 | 3.00E-90 | 100.00% |
| JK507524.1 | 1.00E-89 | 95.85% |
| DR968632.1 | 1.00E-88 | 98.94% |
| CO528041.1 | 2.00E-87 | 99.46% |
| EB816327.1 | 7.00E-87 | 100.00% |
| DR821933.1 | 2.00E-86 | 99.45% |
| DV172761.1 | 9.00E-86 | 98.40% |
| FL135749.1 | 7.00E-82 | 100.00% |
| FL135748.1 | 7.00E-82 | 100.00% |
| FL135745.1 | 7.00E-82 | 100.00% |
| FL135744.1 | 7.00E-82 | 100.00% |
| FL135739.1 | 7.00E-82 | 100.00% |
| EB819544.1 | 7.00E-82 | 100.00% |
| EB401216.1 | 7.00E-82 | 100.00% |
| DY232620.1 | 7.00E-82 | 100.00% |
| CO533759.1 | 7.00E-82 | 100.00% |
| DR828975.1 | 7.00E-82 | 100.00% |
| CD527602.1 | 7.00E-82 | 100.00% |
| AW157956.1 | 7.00E-82 | 100.00% |
| CD441233.1 | 9.00E-81 | 97.31% |
| CD440373.1 | 9.00E-81 | 96.83% |
| FM185777.1 | 3.00E-80 | 99.42% |
| FL135747.1 | 3.00E-80 | 99.42% |
| FL135738.1 | 2.00E-77 | 98.27% |
| CN845482.1 | 7.00E-77 | 98.26% |
| FL135746.1 | 9.00E-76 | 97.66% |
| FL135741.1 | 9.00E-76 | 97.70% |
| FL135737.1 | 4.00E-74 | 97.08% |
| DY534608.1 | 2.00E-73 | 97.62% |
| FL135743.1 | 4.00E-69 | 95.35% |
| FL477397.1 | 3.00E-65 | 98.01% |
| FL435533.1 | 3.00E-65 | 98.01% |
| EE156324.2 | 3.00E-65 | 100.00% |
| EB819543.1 | 3.00E-65 | 100.00% |
| EB165603.1 | 3.00E-65 | 100.00% |
| DY232619.1 | 3.00E-65 | 100.00% |
| CO533758.1 | 3.00E-65 | 100.00% |
| DV543541.1 | 3.00E-65 | 100.00% |
| DV511866.1 | 3.00E-65 | 100.00% |
| DV172760.1 | 3.00E-65 | 100.00% |
| DV029941.1 | 3.00E-65 | 100.00% |
| DR969588.1 | 3.00E-65 | 100.00% |
| DR968631.1 | 3.00E-65 | 100.00% |
| DR964373.1 | 3.00E-65 | 100.00% |
| DR960331.1 | 3.00E-65 | 100.00% |
| DR817073.1 | 3.00E-65 | 100.00% |
| CD527601.1 | 3.00E-65 | 100.00% |
| EB641591.1 | 2.00E-63 | 99.29% |
| DN560261.1 | 7.00E-62 | 98.58% |
| DY401342.1 | 2.00E-58 | 95.36% |
| DV494472.1 | 2.00E-58 | 97.16% |
| CF634947.1 | 2.00E-58 | 97.16% |
| BU037925.1 | 2.00E-58 | 95.36% |
| BQ833798.1 | 2.00E-58 | 95.36% |
| EE010860.1 | 4.00E-54 | 95.07% |
| BQ703828.1 | 7.00E-52 | 94.41% |
| EB401215.1 | 9.00E-51 | 97.58% |
| EB816326.1 | 3.00E-50 | 99.15% |
| CD964905.1 | 7.00E-47 | 95.28% |
| DV523799.1 | 2.00E-33 | 98.85% |

**Table S7.** Stomatal indices and densities of *ali1-1* and wild-type B73 grown in the greenhouse. Data is presented as means with standard deviation of 5 biological replicates measured at V10 stage of the widest portion of leaf 9 of greenhouse grown plants. Letters indicate connecting letter report as determined by Student’s t-test (*P* < 0.05).

| Genotype | SI abaxial<sup>a</sup> | SD abaxial<sup>b</sup> | SI adaxial<sup>a</sup> | SD adaxial<sup>b</sup> | RS<sup>c</sup> |
| --- | --- | --- | --- | --- | --- |
| B73 (+/+) | 20.0 ±1.9 a | 119.3 ±13.5 a | 16.4 ±0.8 a | 72.86 ±5.4 a | 0.82 a |
| *ali1-1/ali1-1* | 18.7 ±1.9 a | 134.3 ±12.5 a | 14.7 ±1.7 a | 73.57 ±5.4 a | 0.79 a |

<sup>a</sup>SI, stomatal index
<sup>b</sup>SD, stomatal density
<sup>c</sup>RS, ratio of stomata adaxial/abaxial

**Table S8.** Stomatal indices and densities of B73, +/*ali1-1*, +/*ali1-2, ali1-1*/*ali1-1, ali1-1*/*ali1-2*, and *ali1-2*/*ali1-2* grown in the field. Data is presented as means with SD of 6 biological replicates measured at V9 stage of the widest portion of leaf 8 of greenhouse grown plants. Letters indicate connecting letter report as determined by ANOVA with posthoc analysis using the Holm-Sidak algorithm (*P* < 0.05).

| Genotype | SI abaxial<sup>a</sup> | SD abaxial<sup>b</sup> |
| --- | --- | --- |
| B73 (+/+) | 24.2 ±1.7 a | 206.5 ±18.8 a |
| *+/ali1-1* | 24.6 ±1.0 a | 212.2 ±12.7 a |
| *+/ali1-2* | 25.1 ±0.5 a | 216.6 ±13.6 a |
| *ali1-1/ali1-1* | 24.2 ±0.7 a | 214.0 ±16.0 a |
| *ali1-1/ali1-2* | 24.1 ±0.6 a | 222.5 ±17.8 a |
| *ali1-2/ali1-2* | 23.7 ±1.4 a | 216.6 ±10.7 a |

<sup>a</sup>SI, stomatal index
<sup>b</sup>SD, stomatal density

**Table S9.** NPC gene expression in *ali1-1* compared to wild type from the tassel node just before emergence from the leaf whorl.

| Gene Name | Maize ID | Mean Exp.<br>(*ali1-1*) | Mean Exp.<br>(wild type) | log2FC | *p*-val. | Genome<br>wide FDR<sup>a</sup> |
| --- | --- | --- | --- | --- | --- | --- |
| ALADIN1 | GRMZM2G180205 | 651.432 | 700.118 | -0.1044 | 0.7105 | 0.8244 |
| RAE1-L1 | GRMZM2G180815 | 1188.697 | 468.662 | 1.3435 | 2.3E-06 | 1.6E-06* |
| RAE1-L2 | GRMZM2G058498 | 2472.891 | 2235.171 | 0.1456 | 0.4998 | 0.6664 |
| CG1 | GRMZM2G089050 | 1131.622 | 593.728 | 0.9296 | 0.0001 | 0.0018* |
| NUP98a | GRMZM2G059015 | 7423.819 | 5070.035 | 0.5500 | 0.0125 | 0.0629 |
| NUP98b | GRMZM2G344924 | 3221.206 | 2210.279 | 0.5431 | 0.0100 | 0.0541 |
| NUP62-L1 | GRMZM2G061043 | 724.381 | 718.645 | 0.0130 | 0.9604 | 0.9778 |
| NUP62-L2 | GRMZM2G162329 | 38.097 | 17.248 | 0.5121 | 0.0905 | 0.7635 |
| NUP214 | AC155434.2_FG004 | 1981.954 | 1277.143 | 0.6325 | 0.0400 | 0.1344 |
| NUP58-L1 | GRMZM2G024775 | 701.122 | 450.021 | 0.6363 | 0.0566 | 0.1663 |
| NUP58-L2 | GRMZM2G134508 | 1990.325 | 1683.229 | 0.2409 | 0.3906 | 0.5701 |
| NUP58-L3 | GRMZM2G345582 | 19.622 | 23.126 | -0.2317 | 0.7876 | 0.8752 |
| NUP54 | GRMZM2G104258 | 1477.189 | 1422.909 | 0.0538 | 0.8163 | 0.8945 |
| NUP136-L1 | GRMZM2G329300 | 2538.364 | 1647.225 | 0.6229 | 0.0308 | 0.1136 |
| NUP136-L2 | GRMZM5G867767 | 160.087 | 63.308 | 1.3265 | 0.1269 | 0.2793 |
| NUP136-L3 | GRMZM5G839512 | 2484.673 | 1423.370 | 0.8024 | 0.0387 | 0.1315 |
| NUP50a | GRMZM2G157317 | 11872.288 | 11422.341 | 0.0559 | 0.7992 | 0.8828 |
| NUP50b | GRMZM5G823017 | 2036.197 | 1906.090 | 0.0961 | 0.6916 | 0.8123 |
| NUP160-L1 | GRMZM2G322493 | 439.143 | 121.458 | 1.8523 | 0.2707 | 0.4518 |
| NUP160-L2 | GRMZM2G167741 | 922.473 | 800.318 | 0.2051 | 0.4184 | 0.5949 |
| NUP133-L1 | GRMZM2G058374 | 1168.932 | 1212.249 | -0.0515 | 0.8279 | 0.9019 |
| NUP133-L2 | GRMZM2G358877 | 489.717 | 221.616 | 1.1426 | 0.4034 | 0.5817 |
| NUP107 | GRMZM2G348675 | 507.382 | 237.073 | 1.0921 | 0.0602 | 0.1731 |
| NUP96-L1 | AC149829.2_FG003 | 494.911 | 525.444 | -0.0847 | 0.7531 | 0.8531 |
| NUP96-L2 | GRMZM5G861959 | 0.653 | 1.585 | -1.3320 | 0.7531 | NA |
| NUP96-L3 | AC207628.4_FG003 | 250.497 | 190.698 | 0.3968 | 0.2334 | 0.4137 |
| NUP75 | GRMZM2G409430 | 1714.633 | 1678.873 | 0.0307 | 0.8932 | 0.9410 |
| NUP43 | GRMZM2G063188 | 1358.505 | 1066.045 | 0.3491 | 0.1250 | 0.2765 |
| SEH1 | GRMZM2G057853 | 2077.787 | 1837.090 | 0.1776 | 0.4015 | 0.5801 |
| SEC13-L1 | GRMZM2G007021 | 5306.593 | 6597.602 | -0.3138 | 0.2197 | 0.3980 |
| SEC13-L2 | GRMZM2G102795 | 37.129 | 30.643 | 0.2650 | 0.7080 | 0.8229 |
| SEC13-L3 | GRMZM2G127850 | 0 | 0 | 0 | NA | NA |
| SEC13-L4 | GRMZM5G871980 | 0 | 0 | 0 | NA | NA |
| SEC13-L5 | GRMZM6G700127 | 9415.549 | 10154.197 | -0.1090 | 0.5966 | 0.7436 |
| SEC13-L6 | GRMZM2G074567 | 3956.764 | 4057.775 | -0.0364 | 0.8580 | 0.9199 |
| NUP205 | GRMZM2G056661 | 2014.200 | 1022.617 | 0.9761 | 0.0069 | 0.0424* |
| NUP188 | GRMZM2G134985 | 1186.793 | 812.309 | 0.5454 | 0.0634 | 0.1791 |
| NUP155 | GRMZM2G077576 | 0 | 0 | 0 | NA | NA |
| NUP93b | GRMZM5G874366 | 0 | 0 | 0 | NA | NA |
| NUP35-L1 | AC187157.4_FG002 | 1364.433 | 1749.168 | -0.3573 | 0.1821 | 0.3530 |
| NUP35-L2 | GRMZM2G084477 | 480.147 | 463.916 | 0.0474 | 0.8934 | 0.9411 |
| NUP88/MOS7 | GRMZM2G156174 | 2394.141 | 1598.607 | 0.5825 | 0.0154 | 0.0717 |
| GP210-L1 | GRMZM2G331533 | 765.072 | 233.323 | 1.7115 | 0.1276 | 0.2805 |
| GP210-L2 | GRMZM2G476810 | 236.117 | 64.308 | 1.8731 | 0.2980 | 0.4803 |
| GP210-L3 | GRMZM2G098622 | 1199.732 | 1004.025 | 0.2570 | 0.2666 | 0.4476 |
| NDC1 | GRMZM2G528010 | 662.933 | 479.523 | 0.4648 | 0.0974 | 0.2358 |
| CPR5-L1 | GRMZM2G032447 | 1307.000 | 876.772 | 0.5744 | 0.0392 | 0.1325 |
| CPR5-L2 | GRMZM2G158811 | 620.545 | 297.668 | 1.0570 | 0.0001 | 0.0024* |
| ELYS/HOS1 | GRMZM2G312738 | 1568.485 | 1176.953 | 0.4136 | 0.0776 | 0.2043 |
| GLE1-L1 | AC210013.4_FG008 | 666.967 | 335.970 | 0.9844 | 0.0096 | 0.0527 |
| GLE1-L2 | GRMZM2G417217 | 498.377 | 280.314 | 0.8252 | 0.0150 | 0.0703 |
| TPR/NUA-L1 | GRMZM2G439339 | 4034.087 | 2716.838 | 0.5706 | 0.0190 | 0.0821 |
| TPR/NUA-L1 | GRMZM2G118403 | 2835.980 | 1968.341 | 0.5261 | 0.0343 | 0.1220 |

<sup>a</sup>Genes that are significantly differentially expressed at a FDR<0.05 is indicated by an asterisk.

**Table S10.** Expression data from tassel tissue of known interactors with the *Homo sapiens* AAAS gene.

Expression data from tassell tissue of known interactors with the *Homo sapiens* AAAS gene.
| Gene Name | Maize ID | *e*-value<sup>a</sup> | Mean Exp.<br>(*ali1-1*) | Mean Exp.<br>(wild type) | log2FC | *p*-val. | Genome<br>wide FDR<sup>b</sup> |
| --- | --- | --- | --- | --- | --- | --- | --- |
| CTDNEP1-L1 | GRMZM2G162347 | 5.0E-36 | 2043.420 | 1471.093 | 0.4738 | 0.0347 | 0.1229 |
| CTDNEP1-L2 | GRMZM2G068690 | 6.7E-36 | 934.978 | 906.122 | 0.0445 | 0.8578 | 0.9198 |
| CTDNEP1-L3 | GRMZM2G122362 | 4.1E-30 | 711.489 | 395.701 | 0.8442 | 0.0017 | 0.0159* |
| CTDNEP1-L4 | GRMZM2G124179 | 1.2E-28 | 809.385 | 950.248 | -0.2304 | 0.3693 | 0.5492 |
| CTDNEP1-L5 | GRMZM2G347623 | 1.7E-28 | 2874.310 | 2159.466 | 0.4120 | 0.0815 | 0.2104 |
| CUL7-L1 | GRMZM2G054247 | 2.1E-05 | 789.579 | 778.669 | 0.0182 | 0.9463 | 0.9705 |
| CUL7-L2 | GRMZM2G174971 | 5.0E-05 | 262.896 | 256.503 | 0.0356 | 0.9097 | 0.9504 |
| EGFR-L1 | GRMZM2G459854 | 6.8E-33 | 899.183 | 956.565 | -0.0891 | 0.6992 | 0.8172 |
| EGFR-L2 | GRMZM2G164242 | 2.5E-32 | 46.117 | 35.624 | 0.3600 | 0.5904 | 0.7388 |
| EGFR-L3 | GRMZM2G104658 | 2.1E-31 | 796.178 | 791.762 | 0.0067 | 0.9783 | 0.9864 |
| EGFR-L4 | GRMZM5G814851 | 3.4E-31 | 416.335 | 331.573 | 0.3259 | 0.2678 | 0.4486 |
| EGFR-L5 | GRMZM2G160922 | 8.5E-31 | 903.125 | 479.683 | 0.9114 | 0.0007 | 0.0084* |
| EGFR-L6 | GRMZM2G152889 | 8.9E-31 | 786.123 | 685.958 | 0.1956 | 0.4417 | 0.6166 |
| EGFR-L7 | GRMZM2G059671 | 7.1E-30 | 1610.035 | 1152.459 | 0.4808 | 0.1362 | 0.2926 |
| EGFR-L8 | GRMZM2G465833 | 1.4E-29 | 1477.344 | 1102.161 | 0.4224 | 0.0656 | 0.1831 |
| EGFR-L9 | GRMZM2G028709 | 1.4E-29 | 349.509 | 225.136 | 0.6304 | 0.0729 | 0.1965 |
| EGFR-L10 | GRMZM2G140612 | 3.0E-27 | 3679.858 | 3997.683 | -0.1195 | 0.6211 | 0.7616 |
| EGFR-L11 | GRMZM2G002531 | 6.2E-27 | 130.670 | 91.644 | 0.5104 | 0.2139 | 0.3917 |
| EGFR-L12 | GRMZM2G102088 | 6.3E-27 | 2345.473 | 1803.048 | 0.3787 | 0.1310 | 0.2849 |
| GNAI3-L1 | GRMZM2G064732 | 6.7E-65 | 1269.870 | 1307.991 | -0.0426 | 0.8468 | 0.9136 |
| GNAI3-L2 | GRMZM2G429113 | 1.5E-26 | 1591.688 | 1210.338 | 0.3948 | 0.1582 | 0.3220 |
| GNAI3-L3 | GRMZM2G016858 | 4.2E-24 | 2492.487 | 1544.867 | 0.6889 | 0.0211 | 0.0885 |
| ISG15-L1 | GRMZM2G116292 | 1.1E-14 | 14409.560 | 17367.044 | -0.2693 | 0.1798 | 0.3505 |
| ISG15-L2 | GRMZM2G118637 | 6.5E-14 | 789.086 | 457.364 | 0.7838 | 0.0059 | 0.0382 |
| ISG15-L3 | GRMZM2G409726 | 6.9E-14 | 89823.173 | 98792.619 | -0.1373 | 0.5664 | 0.7201 |
| ISG15-L4 | GRMZM2G419891 | 6.9E-14 | 71907.360 | 86975.489 | -0.2744 | 0.2363 | 0.4174 |
| KPNB1-L1 | GRMZM2G138419 | 1.2E-168 | 4630.167 | 4926.032 | -0.0894 | 0.6984 | 0.8167 |
| KPNB1-L2 | GRMZM2G066650 | 2.5E-168 | 2405.976 | 2023.540 | 0.2497 | 0.2629 | 0.4440 |
| KPNB1-L3 | GRMZM2G084767 | 8.8E-165 | 5674.799 | 4418.220 | 0.3611 | 0.1123 | 0.2578 |
| KPNB1-L4 | GRMZM2G010452 | 2.6E-154 | 2054.219 | 1486.692 | 0.4658 | 0.0847 | 0.2159 |
| NTRK1-L1 | GRMZM2G059671 | 4.1E-32 | 1610.035 | 1152.459 | 0.4808 | 0.1362 | 0.2926 |
| NTRK1-L2 | GRMZM2G104658 | 2.1E-31 | 796.178 | 791.762 | 0.0067 | 0.9783 | 0.9864 |
| NTRK1-L3 | GRMZM2G098187 | 3.4E-30 | 68.441 | 63.817 | 0.0979 | 0.8521 | 0.9168 |
| NTRK1-L4 | GRMZM2G140612 | 2.5E-29 | 3679.858 | 3997.683 | -0.1195 | 0.6211 | 0.7616 |
| NTRK1-L5 | GRMZM2G102088 | 6.3E-29 | 2345.473 | 1803.048 | 0.3787 | 0.1310 | 0.2849 |
| NTRK1-L6 | GRMZM2G156013 | 1.0E-28 | 4418.520 | 4280.745 | 0.0456 | 0.8326 | 0.9051 |
| NUP107 | GRMZM2G348675 | 1.4E-23 | 507.382 | 237.073 | 1.0921 | 0.0602 | 0.1731 |
| NUP98a | GRMZM2G344924 | 1.1E-26 | 3221.206 | 2210.279 | 0.5431 | 0.0100 | 0.0541 |
| NUP98b | GRMZM2G059015 | 2.6E-26 | 7423.819 | 5070.035 | 0.5500 | 0.0125 | 0.0629 |
| RANBP2-L1 | GRMZM2G162388 | 7.6E-62 | 1.848 | 4.481 | -1.3048 | 0.5340 | NA |
| RANBP2-L2 | GRMZM2G326111 | 8.7E-62 | 184823.555 | 221183.684 | -0.2591 | 0.3017 | 0.4834 |
| RANBP2-L3 | GRMZM2G170397 | 3.6E-57 | 3570.407 | 4783.672 | -0.4217 | 0.0846 | 0.2157 |
| RANBP2-L4 | GRMZM2G063244 | 1.5E-55 | 134.970 | 193.615 | -0.5148 | 0.2181 | 0.3962 |
| RANBP2-L5 | GRMZM2G084521 | 1.5E-54 | 9097.711 | 16212.309 | -0.8334 | 0.0011 | 0.0117* |
| RANBP2-L6 | GRMZM2G087570 | 2.1E-54 | 2461.095 | 3539.270 | -0.5238 | 0.0163 | 0.0742 |
| RANBP2-L7 | GRMZM2G076544 | 1.8E-54 | 1961.014 | 3085.905 | -0.6534 | 0.0075 | 0.0445* |
| RANBP2-L8 | GRMZM2G329306 | 2.1E-50 | 1695.318 | 1624.009 | 0.0614 | 0.7868 | 0.8747 |
| RANBP2-L9 | GRMZM2G401848 | 5.5E-50 | 104.215 | 180.941 | -0.7941 | 0.0499 | 0.1540 |
| RCC1-L1 | GRMZM2G003565 | 1.1E-29 | 2636.331 | 3191.605 | -0.2751 | 0.3210 | 0.5033 |
| RCC1-L2 | GRMZM2G134313 | 8.1E-22 | 619.386 | 448.857 | 0.4630 | 0.1818 | 0.3528 |
| RCC1-L3 | GRMZM2G030542 | 1.2E-20 | 1488.168 | 1270.110 | 0.2287 | 0.3017 | 0.4834 |
| RCC1-L4 | GRMZM2G095562 | 4.2E-20 | 1916.540 | 1804.316 | 0.0868 | 0.6880 | 0.8091 |
| RCC1-L5 | GRMZM2G135770 | 1.3E-19 | 2237.379 | 2500.630 | -0.1612 | 0.5042 | 0.6704 |
| SEH1 | GRMZM2G057853 | 1.2E-33 | 2077.787 | 1837.090 | 0.1775 | 0.4015 | 0.5801 |
| SLC39A4 | GRMZM2G001803 | 5.0E-12 | 2083.655 | 1693.033 | 0.2986 | 0.2388 | 0.4196 |
| UBE2I-L1 | GRMZM2G070047 | 2.4E-73 | 4552.211 | 7914.500 | -0.7977 | 0.0029 | 0.0234* |
| UBE2I-L2 | GRMZM2G063931 | 1.1E-72 | 567.859 | 849.088 | -0.5797 | 0.0466 | 0.1474 |
| UBE2I-L3 | GRMZM2G163398 | 5.0E-72 | 3078.124 | 3305.415 | -0.1029 | 0.6185 | 0.7599 |
| UBE2I-L4 | GRMZM2G312693 | 4.3E-70 | 2421.918 | 2586.496 | -0.0944 | 0.6833 | 0.8056 |
| UBE2I-L5 | GRMZM2G341089 | 4.7E-60 | 185.864 | 287.349 | -0.6254 | 0.0623 | 0.1771 |
| UBE2I-L6 | GRMZM2G038851 | 5.5E-60 | 123.724 | 201.594 | -0.6983 | 0.0895 | 0.2234 |
| UBE2I-L7 | GRMZM2G007260 | 2.7E-34 | 3478.337 | 3976.460 | -0.1927 | 0.3970 | 0.5760 |
| XPO1-L1 | GRMZM2G022258 | 0.0E+00 | 1554.759 | 1327.549 | 0.2277 | 0.2968 | 0.4791 |
| XPO1_L2 | GRMZM2G167031 | 0.0E+00 | 2414.678 | 1803.881 | 0.4212 | 0.0519 | 0.1577 |
| EFNB2 | No Hits | NA | NA | NA | NA | NA | NA |
| IFI16 | No Hits | NA | NA | NA | NA | NA | NA |
| LYPD3 | No Hits | NA | NA | NA | NA | NA | NA |
| NUP153 | No Hits | NA | NA | NA | NA | NA | NA |
| OBSL1 | No Hits | NA | NA | NA | NA | NA | NA |
| POPDC2 | No Hits | NA | NA | NA | NA | NA | NA |
| REL | No Hits | NA | NA | NA | NA | NA | NA |
| TCTN2 | No Hits | NA | NA | NA | NA | NA | NA |
| TCTN3 | No Hits | NA | NA | NA | NA | NA | NA |

<sup>a</sup>e-value score for protein BLAST of *Homo sapiens* AAAS gene interactors against maize genome.
<sup>b</sup>Genes that are significantly differentially expressed at a FDR<0.05 is indicated by an asterisk.

**Table S11.** NPC gene expression in *ali1-1* compared to wild type from SAM and surrounding tissue of 15 DAP plants.

| Gene Name | Maize ID | Mean Exp.<br>(*ali1-1*) | Mean Exp.<br>(wild-type) | log2FC | *p*-val. | Genome<br>wide FDR<sup>a</sup> |
| --- | --- | --- | --- | --- | --- | --- |
| ALADIN1 | GRMZM2G180205 | 85.880 | 86.389 | -0.0134 | 0.9493 | 0.9928 |
| RAE1-L1 | GRMZM2G180815 | 132.748 | 107.678 | 0.3083 | 0.0987 | 0.9928 |
| RAE1-L2 | GRMZM2G058498 | 179.270 | 175.135 | 0.0345 | 0.8262 | 0.9928 |
| CG1 | GRMZM2G089050 | 128.650 | 124.765 | 0.0457 | 0.7986 | 0.9928 |
| NUP98a | GRMZM2G059015 | 1032.689 | 966.776 | 0.0968 | 0.4091 | 0.9928 |
| NUP98b | GRMZM2G344924 | 448.852 | 440.665 | 0.0288 | 0.8402 | 0.9928 |
| NUP62-L1 | GRMZM2G061043 | 80.448 | 91.601 | -0.1937 | 0.4052 | 0.9928 |
| NUP62-L2 | GRMZM2G162329 | 1.322 | 0.905 | 0.5121 | 0.7635 | NA |
| NUP214 | AC155434.2_FG004 | 696.087 | 666.125 | 0.0649 | 0.6729 | 0.9928 |
| NUP58-L1 | GRMZM2G024775 | 83.994 | 84.228 | 0.0077 | 0.9703 | 0.9928 |
| NUP58-L2 | GRMZM2G134508 | 397.645 | 395.540 | 0.0089 | 0.9430 | 0.9928 |
| NUP58-L3 | GRMZM2G345582 | 0 | 0 | 0 | NA | NA |
| NUP54 | GRMZM2G104258 | 255.346 | 245.191 | 0.0565 | 0.6903 | 0.9928 |
| NUP136-L1 | GRMZM2G329300 | 587.544 | 584.512 | 0.0086 | 0.9579 | 0.9928 |
| NUP136-L2 | GRMZM5G867767 | 52.643 | 65.325 | -0.3062 | 0.2465 | 0.9928 |
| NUP136-L3 | GRMZM5G839512 | 677.007 | 752.203 | -0.1522 | 0.2407 | 0.9928 |
| NUP50a | GRMZM2G157317 | 3098.730 | 3250.666 | -0.0689 | 0.5149 | 0.9928 |
| NUP50b | GRMZM5G823017 | 591.142 | 578.312 | 0.0327 | 0.7682 | 0.9928 |
| NUP160-L1 | GRMZM2G322493 | 102.597 | 76.986 | 0.4082 | 0.0584 | 0.9928 |
| NUP160-L2 | GRMZM2G167741 | 99.336 | 84.917 | 0.2262 | 0.2826 | 0.9928 |
| NUP133-L1 | GRMZM2G058374 | 313.158 | 294.686 | 0.0894 | 0.5435 | 0.9928 |
| NUP133-L2 | GRMZM2G358877 | 155.892 | 150.023 | 0.0545 | 0.7620 | 0.9928 |
| NUP107 | GRMZM2G348675 | 98.614 | 75.535 | 0.3843 | 0.1011 | 0.9928 |
| NUP96-L1 | AC149829.2_FG003 | 126.646 | 157.401 | -0.3165 | 0.0679 | 0.9928 |
| NUP96-L2 | GRMZM5G861959 | 0 | 0 | 0 | NA | NA |
| NUP96-L3 | AC207628.4_FG003 | 28.342 | 27.779 | 0.0188 | 0.9507 | 0.9928 |
| NUP75 | GRMZM2G409430 | 261.269 | 266.402 | -0.0309 | 0.8275 | 0.9928 |
| NUP43 | GRMZM2G063188 | 170.235 | 160.168 | 0.0813 | 0.6278 | 0.9928 |
| SEH1 | GRMZM2G057853 | 316.914 | 331.955 | -0.0664 | 0.6431 | 0.9928 |
| SEC13-L1 | GRMZM2G007021 | 777.514 | 816.527 | -0.0709 | 0.4846 | 0.9928 |
| SEC13-L2 | GRMZM2G102795 | 1.022 | 0.287 | 1.7280 | 0.4606 | NA |
| SEC13-L3 | GRMZM2G127850 | 0 | 0 | 0 | NA | NA |
| SEC13-L4 | GRMZM5G871980 | 0 | 0 | 0 | NA | NA |
| SEC13-L5 | GRMZM6G700127 | 2148.770 | 2064.830 | 0.0582 | 0.5255 | 0.9928 |
| SEC13-L6 | GRMZM2G074567 | 665.871 | 710.720 | -0.0941 | 0.3593 | 0.9928 |
| NUP205 | GRMZM2G056661 | 319.553 | 293.978 | 0.1222 | 0.4681 | 0.9928 |
| NUP188 | GRMZM2G134985 | 389.555 | 354.999 | 0.1359 | 0.3537 | 0.9928 |
| NUP155 | GRMZM2G077576 | 0 | 0 | 0 | NA | NA |
| NUP93b | GRMZM5G874366 | 0 | 0 | 0 | NA | NA |
| NUP35-L1 | AC187157.4_FG002 | 235.179 | 244.111 | -0.0532 | 0.7073 | 0.9928 |
| NUP35-L2 | GRMZM2G084477 | 45.326 | 32.765 | 0.4643 | 0.1475 | 0.9928 |
| NUP88/MOS7 | GRMZM2G156174 | 373.742 | 386.135 | -0.0500 | 0.6983 | 0.9928 |
| GP210-L1 | GRMZM2G331533 | 225.572 | 197.771 | 0.1942 | 0.3136 | 0.9928 |
| GP210-L2 | GRMZM2G476810 | 84.224 | 80.400 | 0.0707 | 0.7948 | 0.9928 |
| GP210-L3 | GRMZM2G098622 | 248.775 | 219.502 | 0.1862 | 0.2458 | 0.9928 |
| NDC1 | GRMZM2G528010 | 165.804 | 145.485 | 0.1885 | 0.2767 | 0.9928 |
| CPR5-L1 | GRMZM2G032447 | 345.891 | 380.366 | -0.1351 | 0.3326 | 0.9928 |
| CPR5-L2 | GRMZM2G158811 | 85.185 | 80.471 | 0.0864 | 0.6871 | 0.9928 |
| ELYS/HOS1 | GRMZM2G312738 | 480.723 | 487.181 | -0.0190 | 0.8928 | 0.9928 |
| GLE1-L1 | AC210013.4_FG008 | 154.212 | 163.482 | -0.0902 | 0.6098 | 0.9928 |
| GLE1-L2 | GRMZM2G417217 | 60.343 | 55.063 | 0.1201 | 0.6690 | 0.9928 |
| TPR/NUA-L1 | GRMZM2G439339 | 564.725 | 512.543 | 0.1402 | 0.2931 | 0.9928 |
| TPR/NUA-L1 | GRMZM2G118403 | 1556.200 | 1525.120 | 0.0290 | 0.8387 | 0.9928 |

<sup>a</sup>Genes that are significantly differentially expressed at a FDR<0.05 is indicated by an asterisk.

**Table S12.** Expression data from SAM and surrounding tissue of 15 DAP plants of known interactors with the *Homo sapiens* AAAS gene.

| Gene Name | Maize ID | *e*-value<sup>a</sup> | Mean Exp.<br>(*ali1-1*) | Mean Exp.<br>(wild type) | log2FC | *p*-val. | Genome<br>wide FDR<sup>b</sup> |
| --- | --- | --- | --- | --- | --- | --- | --- |
| CTDNEP1-L1 | GRMZM2G162347 | 5.0E-36 | 398.216 | 452.914 | -0.1855 | 0.1712 | 0.9928 |
| CTDNEP1-L2 | GRMZM2G068690 | 6.7E-36 | 324.369 | 312.716 | 0.0517 | 0.7100 | 0.9928 |
| CTDNEP1-L3 | GRMZM2G122362 | 4.1E-30 | 103.158 | 105.684 | -0.0281 | 0.8951 | 0.9928 |
| CTDNEP1-L4 | GRMZM2G124179 | 1.2E-28 | 75.992 | 77.952 | -0.0425 | 0.8494 | 0.9928 |
| CTDNEP1-L5 | GRMZM2G347623 | 1.7E-28 | 592.198 | 616.130 | -0.0559 | 0.6855 | 0.9928 |
| CUL7-L1 | GRMZM2G054247 | 2.1E-05 | 111.490 | 99.127 | 0.1718 | 0.3864 | 0.9928 |
| CUL7-L2 | GRMZM2G174971 | 5.0E-05 | 15.337 | 18.212 | -0.2342 | 0.6088 | 0.9928 |
| EGFR-L1 | GRMZM2G459854 | 6.8E-33 | 112.242 | 108.420 | 0.0443 | 0.8195 | 0.9928 |
| EGFR-L2 | GRMZM2G164242 | 2.5E-32 | 8.381 | 8.377 | 0.0058 | 0.9916 | 0.9942 |
| EGFR-L3 | GRMZM2G104658 | 2.1E-31 | 72.114 | 82.302 | -0.1831 | 0.4253 | 0.9928 |
| EGFR-L4 | GRMZM5G814851 | 3.4E-31 | 59.450 | 60.970 | -0.0427 | 0.8613 | 0.9928 |
| EGFR-L5 | GRMZM2G160922 | 8.5E-31 | 64.362 | 85.270 | -0.4004 | 0.0773 | 0.9928 |
| EGFR-L6 | GRMZM2G152889 | 8.9E-31 | 18.395 | 25.295 | -0.4498 | 0.2771 | 0.9928 |
| EGFR-L7 | GRMZM2G059671 | 7.1E-30 | 244.731 | 245.410 | -0.0090 | 0.9499 | 0.9928 |
| EGFR-L8 | GRMZM2G465833 | 1.4E-29 | 202.065 | 188.554 | 0.0970 | 0.5394 | 0.9928 |
| EGFR-L9 | GRMZM2G028709 | 1.4E-29 | 80.695 | 62.130 | 0.3638 | 0.1424 | 0.9928 |
| EGFR-L10 | GRMZM2G140612 | 3.0E-27 | 494.580 | 527.403 | -0.0941 | 0.3999 | 0.9928 |
| EGFR-L11 | GRMZM2G002531 | 6.2E-27 | 18.967 | 16.370 | 0.2095 | 0.6340 | 0.9928 |
| EGFR-L12 | GRMZM2G102088 | 6.3E-27 | 346.351 | 304.206 | 0.1891 | 0.1563 | 0.9928 |
| GNAI3-L1 | GRMZM2G064732 | 6.7E-65 | 217.423 | 224.307 | -0.0463 | 0.7604 | 0.9928 |
| GNAI3-L2 | GRMZM2G429113 | 1.5E-26 | 188.580 | 191.180 | -0.0152 | 0.9214 | 0.9928 |
| GNAI3-L3 | GRMZM2G016858 | 4.2E-24 | 437.160 | 460.890 | -0.0732 | 0.5647 | 0.9928 |
| ISG15-L1 | GRMZM2G116292 | 1.1E-14 | 3790.695 | 4405.852 | -0.2171 | 0.0119 | 0.7992 |
| ISG15-L2 | GRMZM2G118637 | 6.5E-14 | 11.371 | 6.424 | 0.8137 | 0.1967 | 0.9928 |
| ISG15-L3 | GRMZM2G409726 | 6.9E-14 | 14699.069 | 14465.199 | 0.0232 | 0.7818 | 0.9928 |
| ISG15-L4 | GRMZM2G419891 | 6.9E-14 | 14048.299 | 13172.773 | 0.0929 | 0.2508 | 0.9928 |
| KPNB1-L1 | GRMZM2G138419 | 1.2E-168 | 833.922 | 770.270 | 0.1156 | 0.2802 | 0.9928 |
| KPNB1-L2 | GRMZM2G066650 | 2.5E-168 | 524.880 | 470.798 | 0.1571 | 0.2606 | 0.9928 |
| KPNB1-L3 | GRMZM2G084767 | 8.8E-165 | 1434.872 | 1311.594 | 0.1299 | 0.2961 | 0.9928 |
| KPNB1-L4 | GRMZM2G010452 | 2.6E-154 | 581.225 | 574.221 | 0.0187 | 0.8933 | 0.9928 |
| NTRK1-L1 | GRMZM2G059671 | 4.1E-32 | 244.731 | 245.410 | -0.0090 | 0.9499 | 0.9928 |
| NTRK1-L2 | GRMZM2G104658 | 2.1E-31 | 72.114 | 82.302 | -0.1831 | 0.4253 | 0.9928 |
| NTRK1-L3 | GRMZM2G098187 | 3.4E-30 | 27.966 | 20.572 | 0.4375 | 0.2496 | 0.9928 |
| NTRK1-L4 | GRMZM2G140612 | 2.5E-29 | 494.580 | 527.403 | -0.0941 | 0.3999 | 0.9928 |
| NTRK1-L5 | GRMZM2G102088 | 6.3E-29 | 346.351 | 304.206 | 0.1891 | 0.1563 | 0.9928 |
| NTRK1-L6 | GRMZM2G156013 | 1.0E-28 | 525.072 | 507.690 | 0.0483 | 0.6644 | 0.9928 |
| NUP107 | GRMZM2G348675 | 1.4E-23 | 98.614 | 75.535 | 0.3843 | 0.1011 | 0.9928 |
| NUP98a | GRMZM2G344924 | 1.1E-26 | 448.852 | 440.665 | 0.0288 | 0.8402 | 0.9928 |
| NUP98b | GRMZM2G059015 | 2.6E-26 | 1032.689 | 966.776 | 0.0968 | 0.4091 | 0.9928 |
| RANBP2-L1 | GRMZM2G162388 | 7.6E-62 | 0.000 | 0.000 | NA | NA | NA |
| RANBP2-L2 | GRMZM2G326111 | 8.7E-62 | 27011.304 | 24675.444 | 0.1305 | 0.2741 | 0.9928 |
| RANBP2-L3 | GRMZM2G170397 | 3.6E-57 | 558.475 | 559.491 | -0.0034 | 0.9810 | 0.9928 |
| RANBP2-L4 | GRMZM2G063244 | 1.5E-55 | 29.214 | 3.030 | 3.2057 | 1.0E-8 | 1.2E-5* |
| RANBP2-L5 | GRMZM2G084521 | 1.5E-54 | 764.391 | 842.285 | -0.1414 | 0.2007 | 0.9928 |
| RANBP2-L6 | GRMZM2G087570 | 2.1E-54 | 87.778 | 78.412 | 0.1621 | 0.4985 | 0.9928 |
| RANBP2-L7 | GRMZM2G076544 | 1.8E-54 | 129.139 | 136.861 | -0.0856 | 0.6552 | 0.9928 |
| RANBP2-L8 | GRMZM2G329306 | 2.1E-50 | 256.530 | 255.727 | -0.0012 | 0.9900 | 0.9940 |
| RANBP2-L9 | GRMZM2G401848 | 5.5E-50 | 26.657 | 24.633 | 0.1294 | 0.7660 | 0.9928 |
| RCC1-L1 | GRMZM2G003565 | 1.1E-29 | 276.773 | 250.599 | 0.1464 | 0.3568 | 0.9928 |
| RCC1-L2 | GRMZM2G134313 | 8.1E-22 | 61.855 | 53.814 | 0.1922 | 0.4553 | 0.9928 |
| RCC1-L3 | GRMZM2G030542 | 1.2E-20 | 180.800 | 215.980 | -0.2577 | 0.0953 | 0.9928 |
| RCC1-L4 | GRMZM2G095562 | 4.2E-20 | 286.505 | 268.633 | 0.0946 | 0.4772 | 0.9928 |
| RCC1-L5 | GRMZM2G135770 | 1.3E-19 | 102.769 | 102.948 | 0.0003 | 0.9551 | 0.9928 |
| SEH1 | GRMZM2G057853 | 1.2E-33 | 316.914 | 331.955 | -0.0664 | 0.6431 | 0.9928 |
| SLC39A4 | GRMZM2G001803 | 5.0E-12 | 154.190 | 170.973 | -0.1509 | 0.3716 | 0.9928 |
| UBE2I-L1 | GRMZM2G070047 | 2.4E-73 | 687.319 | 705.759 | -0.0399 | 0.7139 | 0.9928 |
| UBE2I-L2 | GRMZM2G063931 | 1.1E-72 | 145.395 | 155.929 | -0.1002 | 0.5592 | 0.9928 |
| UBE2I-L3 | GRMZM2G163398 | 5.0E-72 | 560.292 | 617.744 | -0.1427 | 0.2175 | 0.9928 |
| UBE2I-L4 | GRMZM2G312693 | 4.3E-70 | 328.941 | 383.686 | -0.2227 | 0.0775 | 0.9928 |
| UBE2I-L5 | GRMZM2G341089 | 4.7E-60 | 46.832 | 50.660 | -0.1217 | 0.6782 | 0.9928 |
| UBE2I-L6 | GRMZM2G038851 | 5.5E-60 | 2.071 | 3.989 | -0.9445 | 0.3792 | NA |
| UBE2I-L7 | GRMZM2G007260 | 2.7E-34 | 321.966 | 350.489 | -0.1228 | 0.3694 | 0.9928 |
| XPO1-L1 | GRMZM2G022258 | 0.00 | 181.209 | 162.481 | 0.1637 | 0.3514 | 0.9928 |
| XPO1_L2 | GRMZM2G167031 | 0.00 | 337.647 | 285.456 | 0.2434 | 0.0641 | 0.9928 |
| EFNB2 | No Hits | NA | NA | NA | NA | NA | NA |
| IFI16 | No Hits | NA | NA | NA | NA | NA | NA |
| LYPD3 | No Hits | NA | NA | NA | NA | NA | NA |
| NUP153 | No Hits | NA | NA | NA | NA | NA | NA |
| OBSL1 | No Hits | NA | NA | NA | NA | NA | NA |
| POPDC2 | No Hits | NA | NA | NA | NA | NA | NA |
| REL | No Hits | NA | NA | NA | NA | NA | NA |
| TCTN2 | No Hits | NA | NA | NA | NA | NA | NA |
| TCTN3 | No Hits | NA | NA | NA | NA | NA | NA |

<sup>a</sup>e-value score for protein BLAST of Homo sapiens AAAS gene interactors against maize genome.
<sup>b</sup>Genes that are significantly differentially expressed at a FDR<0.05 is indicated by an asterisk.

**Additional Data table S1 (separate file)**

Coding sequence single nucleotide polymorphisms (SNP) between maize inbred B73 and maize inbred Mo17. Column 1 is chromosome number and columns 2 and 3 are SNP positions. (B73_Mo17_SNPs_CDS.bed)

**Additional Data table S2 (separate file)**

DESeq2 results of mRNA differential expression of *ali1-1* tassels before emergence from leaf whorl as compared to wild type. Column 1 is maize identification number, column 2 is the baseMean for all samples, column 3 is the log2 fold change (negative indicates lower expression in *ali1-1* mutant and positive indicates higher expression in *ali1-1* mutant as compared to wild type), column 4 is log2 fold change SE, column 5 is Wald statistic, column 6 is Wald test *P*-value, and column 7 is Benjamini-Hochberg multiple testing adjusted *P*-value. (DESEQ2_ALI_TASSEL.csv)

**Additional Data table S3 (separate file)**

DESeq2 results of mRNA differential expression of *ali1-1* 15 DAP SAM and surrounding tissue as compared to wild type. Column 1 is the maize identification number, column 2 is the baseMean for all samples, column 3 is the log2 fold change (negative indicates lower expression in *ali1-1* mutant and positive indicates higher expression in *ali1-1* mutant as compared to wild type), column 4 is the log2 fold change SE, column 5 is the Wald statistic, column 6 is Wald test *P*-value, and column 7 is the Benjamini-Hochberg multiple testing adjusted *P*-value. (DESEQ2_ALI_15DAP.csv)

**Additional Data table S4 (separate file)**

Filtered list of single nucleotide polymorphisms (SNP) in replicate 1 of *ali1-1* RNA-sequencing mutant pool. Column 1 is chromosome number, column 2 is SNP position, column 3 is reference nucleotide, column 4 is non-reference nucleotide, column 5 is forward strand reference read count, column 6 is reverse strand reference read count, column 7 is forward strand non-reference read count, column 8 is reverse strand non-reference read count, column 9 is snpEff annotation of the SNP, column 10 is highest Arabidopsis BLASTP hit, column 11 is e-value score for BLAST, and column 12 is Arabidopsis annotation. (ALI1_REP1_RNA_CALLSNPs.tsv)

**Additional Data table S5 (separate file)**

Filtered list of single nucleotide polymorphisms (SNP) in replicate 2 of *ali1-1* RNA-sequencing mutant pool. Column 1 is chromosome number, column 2 is SNP position, column 3 is reference nucleotide, column 4 is non-reference nucleotide, column 5 is forward strand reference read count, column 6 is reverse strand reference read count, column 7 is forward strand non-reference read count, column 8 is reverse strand non-reference read count, column 9 is snpEff annotation of the SNP, column 10 is highest Arabidopsis BLASTP hit, column 11 is e-value score for BLAST, and column 12 is Arabidopsis annotation. (ALI1_REP2_RNA_CALLSNPs.tsv)

**Additional Data table S6 (separate file)**

Filtered list of single nucleotide polymorphisms (SNP) in replicate 1 of wild-type RNA-sequencing pool. Column 1 is chromosome number, column 2 is SNP position, column 3 is reference nucleotide, column 4 is non-reference nucleotide, column 5 is forward strand reference read count, column 6 is reverse strand reference read count, column 7 is forward strand non-reference read count, column 8 is reverse strand non-reference read count, column 9 is snpEff annotation of the SNP, column 10 is highest Arabidopsis BLASTP hit, column 11 is e-value score for BLAST, and column 12 is Arabidopsis annotation. (WT_REP1_RNA_CALLSNPs.tsv)

**Additional Data table S7 (separate file)**

Filtered list of single nucleotide polymorphisms (SNP) in replicate 2 of wild-type RNA-sequencing pool. Column 1 is chromosome number, column 2 is SNP position, column 3 is reference nucleotide, column 4 is non-reference nucleotide, column 5 is forward strand reference read count, column 6 is reverse strand reference read count, column 7 is forward strand non-reference read count, column 8 is reverse strand non-reference read count, column 9 is snpEff annotation of the SNP, column 10 is highest Arabidopsis BLASTP hit, column 11 is e-value score for BLAST, and column 12 is Arabidopsis annotation. (WT_REP2_RNA_CALLSNPs.tsv)

**Additional Data table S8 (separate file)**

Filtered list of single nucleotide polymorphisms (SNP) in 9 individual *ali1-1*; BC1 F2 plants of DNA-sequencing. Column 1 is chromosome number, column 2 is SNP position, column 3 is reference nucleotide, column 4 is non-reference nucleotide, column 5 is forward strand reference read count, column 6 is reverse strand reference read count, column 7 is forward strand non-reference read count, column 8 is reverse strand non-reference read count, column 9 is snpEff annotation of the SNP, column 10 is highest Arabidopsis BLASTP hit, column 11 is e-value score for BLAST, and column 12 is Arabidopsis annotation. (ALI1_DNA_CALLSNPs.zip)

**Additional Data table S9 (separate file)**

Selected gene list of DESeq2 results of mRNA differential expression of *ali1-1* tassels before emergence from leaf whorl as compared to wild type. Column 1 is maize identification number, column 2 is the baseMean for all samples, column 3 is the log2 fold change (negative indicates lower expression in *ali1-1* mutant and positive indicates higher expression in *ali1-1* mutant as compared to wild type), column 4 is log2 fold change SE, column 5 is Wald statistic, column 6 is Wald test *P*-value, and column 7 is Benjamini-Hochberg multiple testing adjusted *P*-value. (INTEREST_TASSEL.csv)

**Additional Data table S10 (separate file)**

Selected gene list of DESeq2 results of mRNA differential expression of *ali1-1* 15 DAP SAM and surrounding tissue as compared to wild type. Column 1 is the maize identification number, column 2 is the baseMean for all samples, column 3 is the log2 fold change (negative indicates lower expression in *ali1-1* mutant and positive indicates higher expression in *ali1-1* mutant as compared to wild type), column 4 is the log2 fold change SE, column 5 is the Wald statistic, column 6 is Wald test *P*-value, and column 7 is the Benjamini-Hochberg multiple testing adjusted *P*-value. (INTEREST_15DAP.csv)

